# A fluorophore–ligand conjugate for the rapid and reversible staining of native AMPA receptors in living neurons

**DOI:** 10.1101/2024.08.21.608930

**Authors:** Kyohei Soga, Takaaki Fujiwara, Mayu Nakagawa, Akihiro Shibata, Hansel Adriel, Kenji Yatsuzuka, Wataru Kakegawa, Michisuke Yuzaki, Itaru Hamachi, Eriko Nango, Shigeki Kiyonaka

## Abstract

The subcellular localizations of neurotransmitter receptors are strictly regulated in neurons. Changes in the trafficking of α-amino-3-hydroxy-5-methyl-4-isoxazole-propionic acid (AMPA)-type glutamate receptors (AMPARs) play an essential role in synaptic plasticity, which is the cellular basis of learning and memory. To explore receptor trafficking, genetically encoded approaches (e.g., the fusion of fluorescent proteins to receptors) are often used. However, concerns remain that genetic approaches cannot fully reproduce the receptor function that is inherent to neurons. Herein, we report a chemical probe, PFQX1(AF488), for the visualization of cell-surface AMPARs without any genetic manipulation to neurons. The rapid and reversible staining features of this probe allowed the visualization of AMPAR accumulation in dendritic spines during synaptic plasticity in living hippocampal neurons. Moreover, the mechanisms of this synaptic accumulation, for which genetically encoded approaches have given controversial results, were revealed by integrating two chemical methods: PFQX1(AF488) and covalent chemical labeling.

## Introduction

Synapses are the fundamental units of brain function, and their structure and function are strengthened or weakened over time in response to a variety of inputs. This phenomenon is known as synaptic plasticity, which is a fundamental process that underlies learning and memory acquisition^1–3^. In particular, excitatory postsynaptic currents mediated by ionotropic glutamate receptors are enhanced by repeated synaptic activity; this is known as long-term potentiation (LTP).

α-Amino-3-hydroxy-5-methyl-4-isoxazole-propionic acid (AMPA)-type glutamate receptors (AMPARs) are the most abundant ionotropic glutamate receptors. They play essential roles in learning and memory^4,5^. AMPARs are localized not only on the postsynaptic terminal but also intracellularly and in extrasynaptic regions such as the dendritic surface. The subcellular localization of AMPARs is dynamically regulated by endocytosis, exocytosis, and lateral diffusion^4–6^. Many studies have suggested that changes in AMPAR trafficking may have essential roles in the expression of LTP^7–9^. However, further clarification of these mechanisms is necessary to better understand the molecular mechanisms of memory.

To analyze AMPAR trafficking, fluorescent proteins can be fused to the extracellular domain of AMPARs using genetic approaches. Notably, a pH-sensitive variant of green fluorescent protein (super-ecliptic pHluorin [SEP]) is widely used for visualizing cell-surface AMPARs in neurons^7,8,10,11^. Self-labeling protein tags such as SNAP-^12^ or Halo-tags^13,14^ with cell-impermeable fluorescent probes have also been used instead of fluorescent proteins. Importantly, although these protein tags are undoubtedly powerful for studying AMPAR trafficking in neurons, they largely rely on the overexpression of genetically encoded receptors. Given that the number of AMPAR is strictly controlled at synapses^4,15^, their expression levels, post-translational modifications, and interactions with accessory proteins are likely to be affected by the overexpression of tagged AMPARs. To resolve this critical problem, a knock-in strategy has been reported^16,17^ in which SEP is inserted into the N-terminal region of the target AMPAR subunit gene on the genome. However, even in knock-in mice, the expression levels of target AMPAR are affected. Furthermore, considering that the N-terminal region of ionotropic glutamate receptors interacts with synaptic proteins, the introduction of a large protein tag to this region may affect AMPAR function. Similarly, the introduction of the exogenous gene into the genome itself may affect the transcription and/or stability of AMPAR mRNA^16^.

Ideally, native AMPARs would be analyzed to clarify their physiological roles without any genetic manipulation. In this context, we have previously reported the fluorescent labeling of cell-surface AMPARs endogenously expressed in neurons using ligand-directed acyl imidazole chemistry^18–20^. In this method, a fluorescent dye-bearing chemical reagent, termed chemical AMPAR modification 2 (CAM2), is recognized by AMPARs, and amino acid residues (e.g., Lys or Ser) near the ligand-binding domain (LBD) of AMPARs are covalently labeled with the fluorophore (**Extended Data Fig. 1a**). Although the original CAM2 method is powerful for visualizing AMPARs not only in cultured neurons^19^ but also in vivo^20^, it is associated with some restrictions, such as a long incubation period (1–4 h) at a low temperature (e.g., 17 °C) to suppress the internalization of fluorophore-labeled receptors in cultured neurons. To overcome these limitations, we have reported an alternative method, termed ligand-directed two-step labeling, in which ligand-directed acyl imidazole chemistry is combined with click chemistry (**Extended Data Fig. 1b**)^21^. Although a relatively long incubation period at 37 °C is still needed for labeling a *trans*-cyclooctene (TCO) to AMPARs in the first step, the fluorophore is covalently labeled to TCO-labeled AMPARs on the cell surface within 5 min via a fast click chemistry reaction in the second step. This method is powerful for analyzing the fate of cell-surface AMPARs. However, to analyze the trafficking of AMPARs during synaptic plasticity, both the cell surface and exocytosed components need to be visualized and quantified, and it remains challenging to analyze intracellular AMPARs using these methods. Thus, a new method that can analyze intracellular AMPARs is required.

In the present study, we report a fluorophore–ligand conjugate, termed PFQX1(AF488), for the rapid and reversible visualization of AMPARs in neurons. This probe allowed us to stain cell-surface AMPARs immediately after its addition to the medium. Thanks to its reversible and reproducible staining features, PFQX1(AF488) successfully visualized the trafficking of AMPARs, including the exocytosis step, during LTP. Furthermore, we were able to elucidate the mechanisms of this trafficking by combining PFQX1(AF488) with ligand-directed chemical labeling.

## Results

### Development of fluorophore–ligand conjugates for visualizing cell-surface AMPARs

To visualize AMPARs on the cell surface, we designed cell-impermeable fluorescent probes that were capable of binding to AMPARs (**Fig. 1a, b**). For selective binding to AMPARs, we focused on an anionic ligand, 6-pyrrolyl-7-trifluoromethyl-quinoxaline-2,3-dione (PFQX), which has been developed as a selective antagonist for AMPARs among glutamate receptors, with a high affinity (*K_i_* = 170 nM)^22^. Fluorescein (Fl), which has anionic, hydrophilic, and structurally compact features, was selected as the fluorophore. Next, Fl was fused to the pyrrole moiety of PFQX according to the structural information of AMPARs with ZK200775^23^, a PFQX analog. We synthesized both PFQX1(Fl), in which PFQX and Fl were directly connected, and PFQX2(Fl), which contained an ethylene glycol linker between PFQX and Fl (**Fig. 1b** and **Extended Data Fig. 2**).

**Fig. 1.**
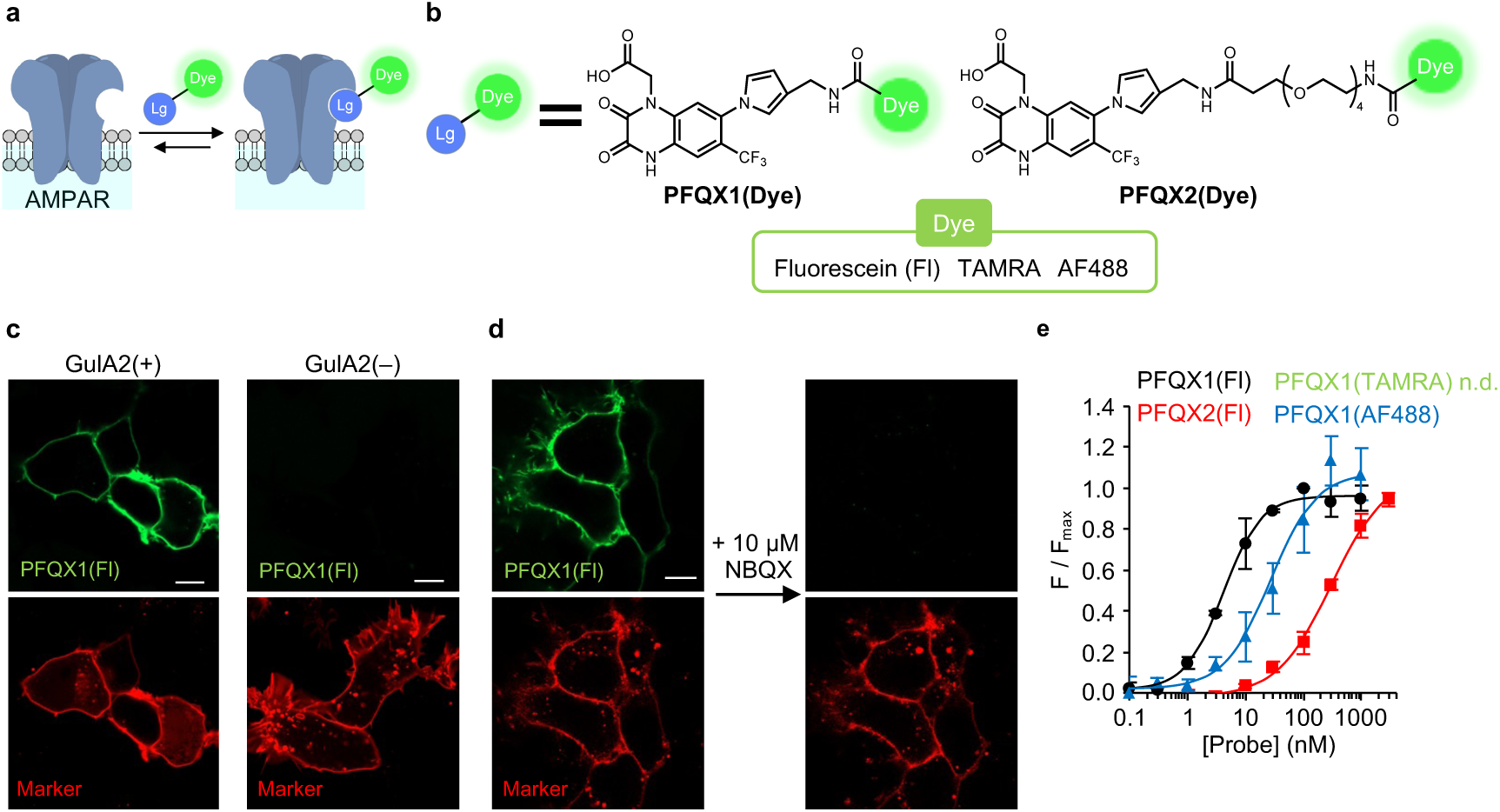
Development of fluorophore–ligand conjugates for AMPARs. **(a)** Schematic illustration of AMPAR staining with fluorophore–ligand conjugates. Lg; AMPAR ligand, Dye; fluorescent dye. **(b)** Chemical structures of fluorophore–ligand conjugates. See Extended Data Fig. 2 for detailed chemical structures. **(c)** Confocal live images of HEK293T cells transfected with GluA2 (GluA2(+)) or its control vector (GluA2(–)). The cells were treated with 100 nM PFQX1(Fl). mCheey-F was used as a transfection marker. Scale bars, 10 µm. **(d)** Competitive inhibition of PFQX1(Fl) binding by NBQX. 100 nM PFQX1(Fl) was added to the HEK293T cells expressed with GluA2. Then, 10 µM NBQX was added to the medium. Scale bar, 10 µm. **(e)** Concentration-dependent binding of fluorophore–ligand conjugates to GluA2 (n = 3). See Extended Data Fig. 3a for detailed results. Data are represented as mean ± s.e.m.

AMPARs form tetrameric cation channels that are composed of GluA1–GluA4 subunits^24^, of which GluA2 is the most abundant^25^. Using confocal microscopy, we then analyzed the binding of the synthesized probes to AMPARs in HEK293T cells that transiently expressed GluA2. After the addition of 100 nM PFQX1(Fl) to the culture dish, prominent fluorescent signals were observed from the cell surface of GluA2-transfected cells but not control cells (**Fig. 1c**). Importantly, the fluorescent signals disappeared with the addition of 10 μM NBQX, a competitive ligand (**Fig. 1d**), suggesting that the fluorescent signals were driven by ligand recognition. As shown in Fig. 1e, the fluorescent signals occurred in a probe concentration-dependent manner. Using PFQX1(Fl), cell-surface fluorescence was clearly observed at a probe concentration of 3 nM, and the signals were saturated at 30 nM of probe (**Fig. 1e** and **Extended Data Fig. 3a**). However, using PFQX2(Fl), which had an ethylene glycol linker, the addition of 30 nM of probe led to only a weak fluorescent signal from the cell surface, and prominent signals were obtained at concentrations of 100 nM and greater. Although the fluorescent signals of PFQX2(Fl) were saturated at 1,000 nM, fluorescence became visible even in the medium. The binding affinity of fluorophore–ligand conjugates with GluA2 was then determined from the cell-surface signal; the *K_d_* values for PFQX1(Fl) and PFQX2(Fl) were 4.3 ± 0.5 nM and 346 ± 99 nM, respectively (**Fig. 1e**). Interestingly, the affinity of PFQX1(Fl) was higher than those reported for both PFQX (*K_i_* = 170 nM)^22^ and PFQX-amine (*K_i_* = 477 ± 171 nM), the synthetic intermediate of PFQX1(Fl) (**Extended Data Fig. 3b**). This finding indicates that the fluorophore moiety of PFQX1(Fl) may contribute to increasing its affinity to GluA2.

To further explore the effects of the fluorophore on PFQX1(Dye) affinity, we synthesized PFQX1(TAMRA) and PFQX1(AF488) (**Fig. 1b** and **Extended Data Fig. 2**). The fluorescent visualization of AMPARs using these probes was evaluated in HEK293T cells, as described in the previous paragraph. With PFQX1(TAMRA), which had a cationic rhodamine fluorophore, only weak cell-surface fluorescence was visualized, even at a probe concentration of 100 nM (**Extended Data Fig. 3a**). Although the cell-surface fluorescent signal corresponding to AMPARs became more visible at 1,000 nM, the fluorescent intensity from the extracellular solution was the same level as that from the cell surface. Conversely, with PFQX1(AF488), which had a rhodamine backbone but two sulfonate groups, cell-surface fluorescence was observed upon the addition of relatively low probe concentrations (10 nM). The *K_d_* value of PFQX1(AF488) to GluA2 in HEK293T cells was 37.2 ± 9.8 nM, indicating that the anionic xanthene dye in PFQX1(Dye) would contribute to increasing the affinity to AMPARs (**Fig. 1e** and **Extended Data Fig. 3a**).

Given that neuronal activity may change pH homeostasis^26^, we next tested the pH dependency of the fluorescent intensity of the two high-affinity probes (PFQX1(Fl) and PFQX1(AF488)) to AMPARs. As shown in **Extended Data Fig. 4**, the fluorescent intensity of PFQX1(Fl) was weakened even at mild acidic conditions (e.g., pH 6), whereas that of PFQX1(AF488) was relatively unaffected at pH 5–10. Thus, PFQX1(AF488) likely allows the quantification of cell-surface AMPARs without being affected by pH changes in extracellular spaces.

### Structural analysis of the fluorophore–ligand conjugate with AMPARs

As described in the previous section, the affinity of PFQX1(AF488) to AMPARs is one order higher than that of the original ligand PFQX. To elucidate the binding manner of PFQX1(AF488), we conducted structural analyses of AMPAR bound to PFQX1(AF488). AMPARs form tetrameric ion channels, and each subunit is composed of an amino-terminal domain, a ligand-binding domain (LBD), and a transmembrane domain (**Extended Data Fig. 5a**)^24^. Competitive antagonists, including PFQX, bind to the LBD^19,22^. We thus used a ligand-binding core moiety of GluA2 LBD, termed S1S2J^27,28^, for the X-ray crystal structural analysis, and the structure of the S1S2J bound to PFQX1(AF488) was determined at 1.82 Å (**Fig. 2a** and **Supplementary Table 1**). In the structural representation, the residue numbers of S1S2J are shown; the corresponding residue of full-length GluA2 is also shown in parentheses.

**Fig. 2.**
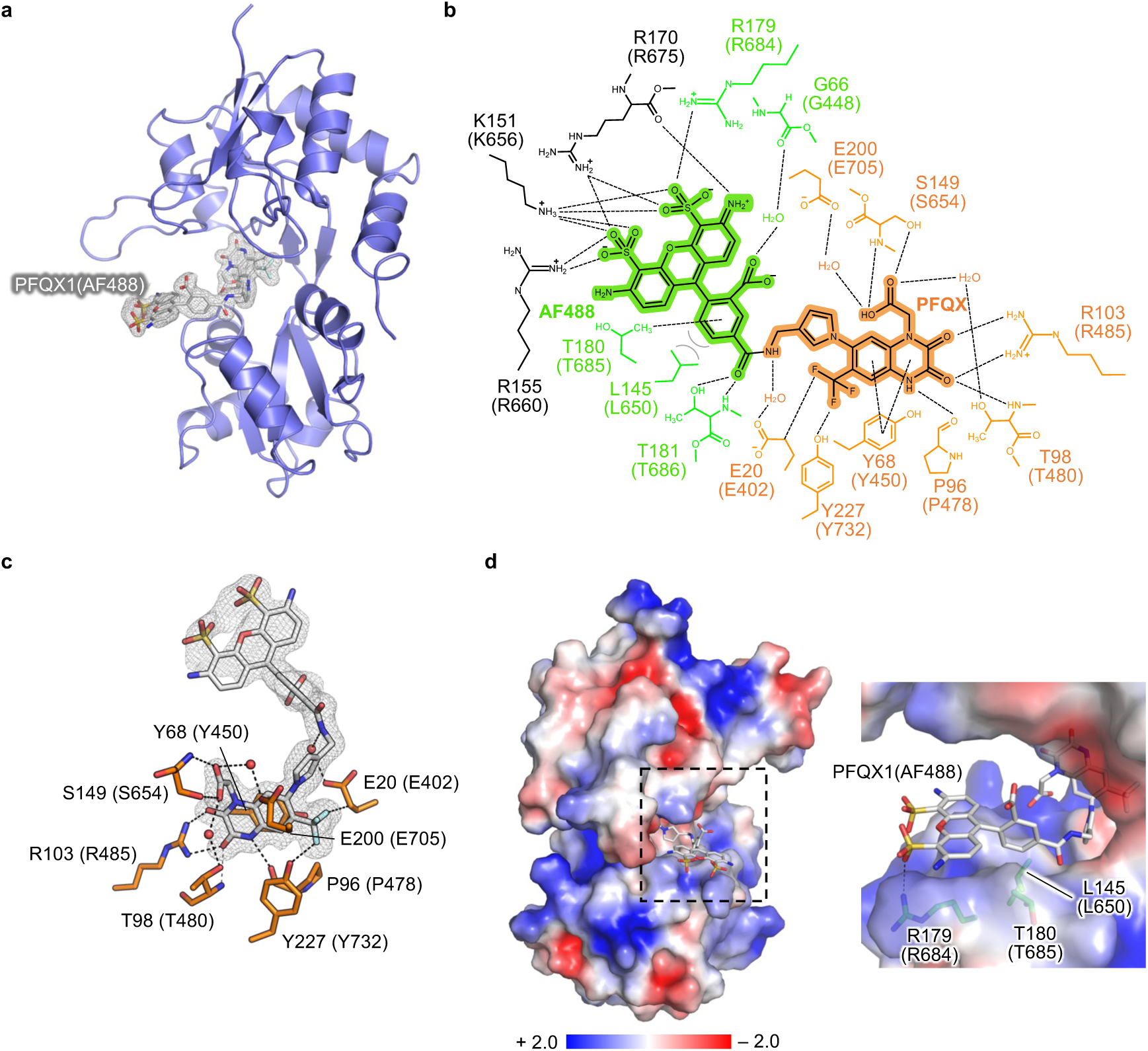
Structure of the S1S2J complexed with PFQX1(AF488). The residue numbers of S1S2J are shown, and the corresponding residue of full-length GluA2 is also shown in parentheses. **(a)** The overall structure of the S1S2J bound to PFQX1(AF488) is displayed as a ribbon diagram. The electron density of bound PFQX1(AF488) (gray) was shown as the polder map at contoured level of 3.0σ. **(b)** Interaction manner of the PFQX1(AF488). The PFQX and AF488 parts of the PFQX1(AF488) are highlighted in orange and green, respectively. The residues and waters contacted to the PFQX and AF488 are also shown in orange and green. The interacting residues in the symmetry mate are coloured in black. Polar and hydrophobic interactions are shown as dotted lines (black) and solid lines (gray), respectively. The atoms in the PFQX1(AF488) are assigned in the upper right panel. **(c)** Contacts between the PFQX part and S1S2J. The residues interacted with the PFQX part are shown as sticks (orange). The polder map of the PFQX1(AF488) is shown at 3.0σ. **(d)** The APBS-generated electrostatic potential of the S1S2J. A positively charged surface around the AF488 is extended. Right, close-up view of the ligand binding site, and the residues involved in recognizing the AF488 part are shown as sticks (green).

Many studies have reported that GluA2 LBD and S1S2J maintain an open form when the antagonist binds to the hinge region of its clamshell-like architecture^28,29^. In the obtained S1S2J-PFQX1(AF488) structure, the S1S2J region showed an open form, which was almost identical to the previously reported structure bound with NBQX^30^ (a well-known antagonist), with a root mean square deviation of 0.64 Å for 258 C^α^ atoms (**Extended Data Fig. 5b**). In the structure, PFQX1(AF488) was recognized over a large area of the aperture of S1S2J (**Fig. 2b** and **Supplementary Table 2**).

The PFQX part of PFQX1(AF488) was deeply bound to the aperture of the S1S2J and showed a similar binding mode to that of ZK200775^31^, which is a structural analog of PFQX (**Fig. 2c** and **Extended Data Fig. 5c**). Specifically, the aromatic group of PFQX was stacked with Tyr68(Tyr450). The two carbonyl groups of PFQX, which mimic the α-carboxyl group of glutamate, formed hydrogen bonds with Arg103(Arg485) and Thr98(Thr480). In contrast to the PFQX part, the AF488 fluorophore part of PFQX1(AF488) was located at the vestibule of the binding cleft of S1S2J. In addition, the phenyl group of AF488 was situated at the hydrophobic region surrounded by Leu145(Leu650) and Thr180(Thr685). Thr180(Thr685) formed the CH-π interaction with the phenyl group. The main-chain amide group as well as the side chain of Thr181(Thr686) formed a hydrogen bond with the carbonyl group of the amide group of the AF488 part (**Fig. 2b**). Of note, the negatively charged sulfonate group of AF488 interacted with the side chain of Arg179(Arg684) (**Fig. 2d**). In addition, the side chain of Lys67(Lys449) and the main-chain amide bonds consisting of Asp146(Asp651)– Ser147(Ser652)–Gly148(Gly653) (referred to as the DSG loop) provided an electropositive environment that was favorable to locating the sulfonate group of AF488 (**Fig. 2d** and **Extended Data Fig. 5d**). Thus, the X-ray structural analysis clearly revealed the molecular basis of the higher affinity of PFQX1(AF488) bearing an anionic fluorophore for AMPARs.

### Rapid and reversible staining of cell-surface AMPARs in HEK293T cells

After developing PFQX1(AF488), a high-affinity probe for AMPARs, we applied this probe for the fluorescent imaging of AMPAR in living cells. To quantitatively analyze the amount of cell-surface AMPARs in each period during live imaging, this fluorescent staining needed to occur in a rapid, reversible, and reproducible manner. We therefore analyzed the kinetics of PFQX1(AF488) binding to AMPARs in HEK293T cells expressed with GluA2 (**Fig. 3a**). As shown in Fig. 3b, prominent fluorescence was observed from the cell surface just 10 s after the addition of 100 nM PFQX1(AF488) to the culture dish. This fluorescent intensity was saturated within 20 s after the addition of the probe, thus demonstrating its rapid binding kinetics.

**Fig. 3.**
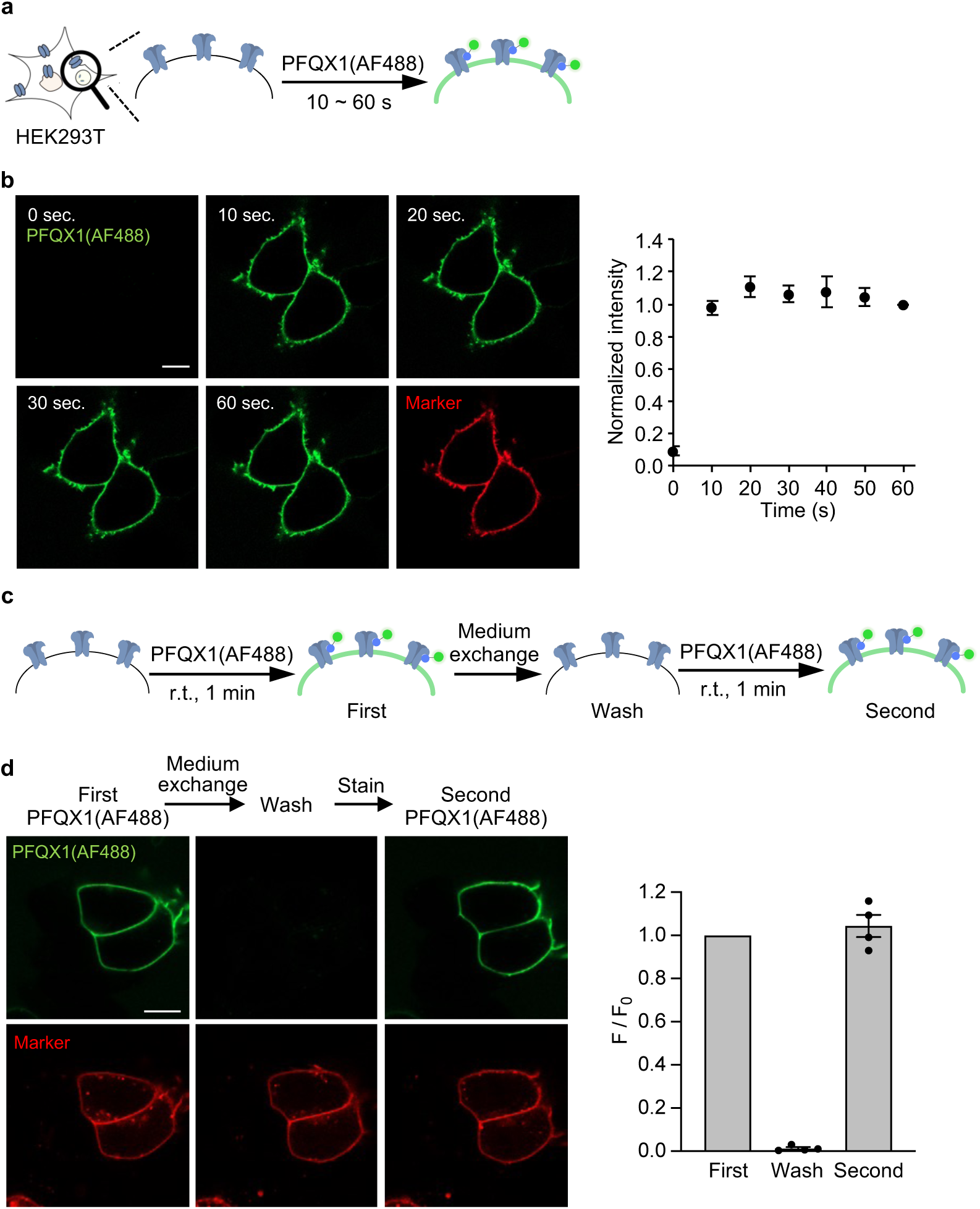
Rapid and reversible staining of AMPARs in HEK293T cells. **(a)** The experimental procedure to analyze the time-course of PFQX1(AF488) binding to GluA2 on live cells. The fluorescent images were obtained at specified time points after the addition of 100 nM PFQX1(AF488) by confocal microscopy. **(b)** Left, representative confocal live images of HEK293T cells expressing Halo-tag-fused GluA2 on the N-terminus after the addition of 100 nM PFQX1(AF488). Halo-tag ligand AF647 was used as a transfection marker. Right, the surface intensity of AF488 was quantified, which was normalized by the intensity at 60 sec. (n = 3). Scale bar, 10 µm. **(c)** The experimental procedure to analyze the reversibility of PFQX1(AF488) binding to GluA2. First image was obtained 1 min after addition of 100 nM PFQX1(AF488) to the culture medium. A wash image was obtained after the medium exchange. Then, 100 nM PFQX1(AF488) was re-treated to obtain the second image. **(d)** Left, representative confocal live cell imaging of HEK293T cells expressed with GluA2 to analyze the reversibility of PFQX1(AF488) binding. Right, the surface intensity of AF488 was quantified, which was normalized by the first images (n = 4). Scale bar, 10 µm. Data are represented as mean ± s.e.m.

To evaluate reversibility, after AMPARs were visualized using 100 nM PFQX1(AF488), the washing out of bound PFQX1(AF488) was examined via medium exchange (**Fig. 3c**). Subsequently, the same concentration of probe was again added to the dish to confirm reproducibility. As shown in Fig. 3d, fluorescence from the cell surface was almost lost after the medium exchange in the same region of view, thus demonstrating the reversibility of PFQX1(AF488) binding to AMPARs. When 100 nM PFQX1(AF488) was again added to the dish, prominent fluorescence from the cell surface was observed at the same level as that obtained after the first treatment. Together, these results indicate that cell-surface AMPAR levels can be quantified repeatedly, even after exchanging the culture medium.

AMPARs form tetrameric ion channels composed of GluA1–4 subunits^24^, and PFQX can antagonize AMPARs in a subunit-nonspecific manner (**Extended Data Fig. 6a**). Consistently, the addition of PFQX1(AF488) led to the visualization of not only GluA2 but also GluA1, both of which are highly expressed in hippocampal neurons^25^ (**Extended Data Fig. 6b, c**). This finding indicates that the probe can visualize AMPARs regardless of subunit type.

### Visualization of endogenous AMPARs in living hippocampal neurons

We next evaluated PFQX1(AF488) for the visualization of endogenous AMPARs in primary cultured hippocampal neurons. In the cultured neurons, a cytosolic marker that is applicable to live cell imaging (CellTracker Red or Calcein Red-AM) was added to the culture medium to stain the intracellular region of the neurons. After the addition of 100 nM of PFQX1(AF488), fluorescent puncta of PFQX1(AF488) were observed along dendrites in the hippocampal neurons pre-treated with the cytosolic marker (**Fig. 4a, b**). Immunostaining for postsynaptic density 95 (PSD95) gave similar punctate signals along dendrites (**Extended Data Fig. 7a**), suggesting that the PFQX1(AF488) puncta correspond to dendritic spines in the neurons. Furthermore, the fluorescent signals from the puncta almost disappeared with the addition of 10 μM NBQX, a competitive ligand (**Fig. 4c**); this finding further indicates that the fluorescent puncta correspond to AMPAR signals.

**Fig. 4.**
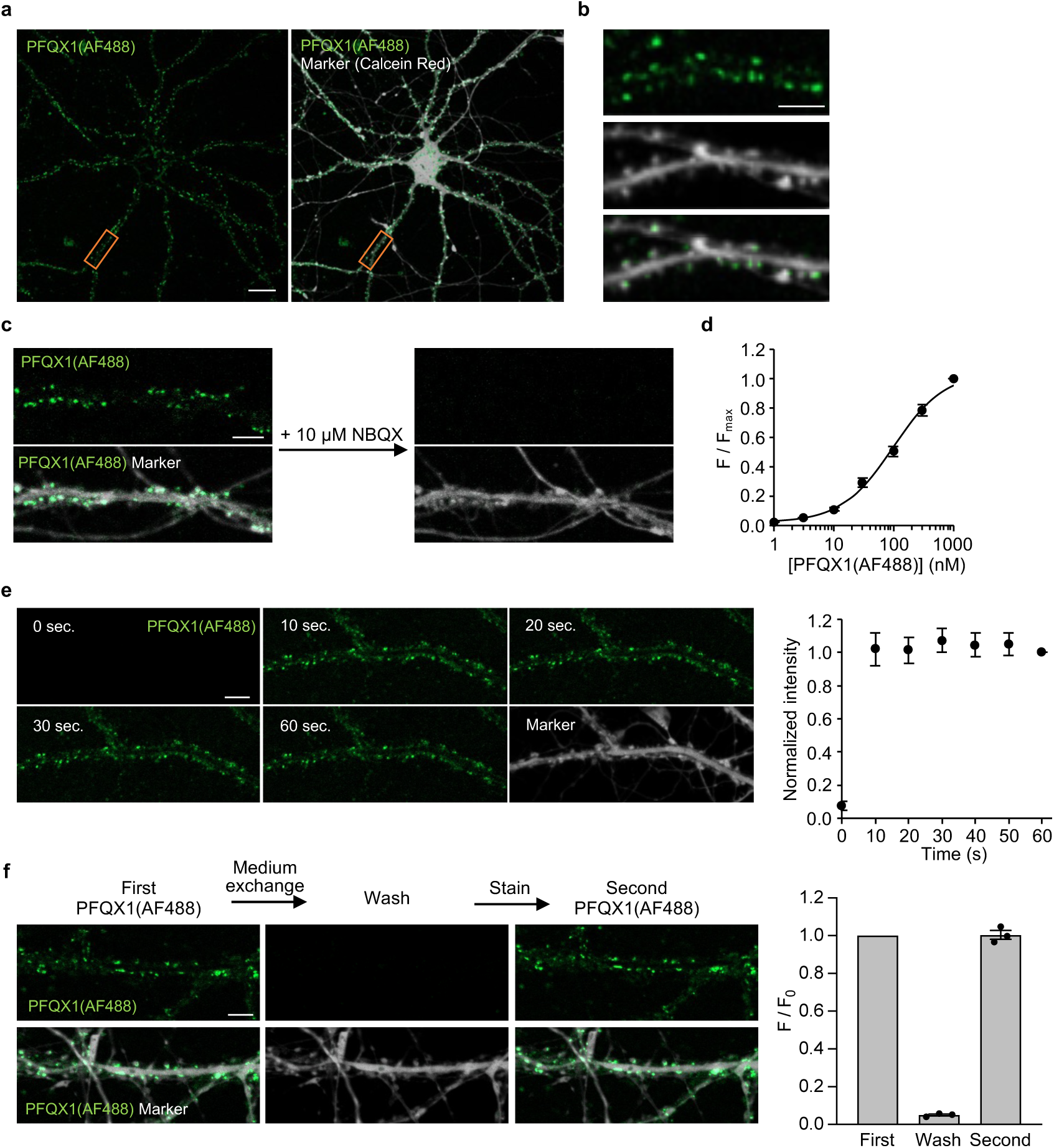
Visualization of native AMPARs in cultured hippocampal neurons. **(a, b)** Confocal live images of cultured hippocampal neurons treated with 100 nM PFQX1(AF488). Calcein red signals are shown in gray as a cytoplasmic marker. Orange ROIs indicated in **(a)** are expanding in **(b)**. Scale bars, 20 µm in **a** and 5 µm in **b**. **(c)** Competitive inhibition of PFQX1(AF488) binding by NBQX. 100 nM PFQX1(AF488) was added to the cultured hippocampal neurons. Then, 10 µM NBQX was added to the medium. Scale bar, 5 µm. **(d)** Concentration-dependency of PFQX1(AF488) binding to AMPARs in cultured hippocampal neurons (n = 3). **(e)** Time-course of PFQX1(AF488) binding. The cells were treated with 100 nM PFQX1(AF488), and images were obtained at specified time points by confocal microscopy. Left, representative confocal live images of cultured hippocampal neurons after the addition of 100 nM PFQX1(AF488). Right, the intensity of PFQX1(AF488) in spines was quantified, which was normalized by the intensity at 60 sec. (n = 3). Scale bar, 5 µm. **(f)** Reversibility of PFQX1(AF488) binding to AMPARs. Left, representative confocal live imaging of cultured hippocampal neurons. Right, the intensity of PFQX1(AF488) in spines was quantified, which was normalized by the first images. [PFQX1(AF488)] = 100 nM (n = 3). Scale bar, 5 µm. Data are represented as mean ± s.e.m.

We next conducted a more detailed investigation into the binding properties of PFQX1(AF488) to AMPARs in cultured hippocampal neurons. As shown in **Extended Data Fig. 7b**, fluorescent signals of PFQX1(AF488) were observed in a concentration-dependent manner. The *K_d_* value of PFQX1(AF488) was 97.5 ± 9.9 nM (**Fig. 4d**) in neurons, which was slightly higher but generally comparable with that obtained in HEK293T cells transfected with AMPARs (**Extended Data Fig. 6c**). As was the case in HEK293T cells, the binding kinetics of PFQX1(AF488) to AMPARs were rapid, and the fluorescent signals were saturated after 10 s of probe addition to the medium (**Fig. 4e**). As shown in Fig. 4f, the fluorescence was easily removed by medium exchange, and re-treatment with PFQX1(AF488) produced fluorescent intensity at the same level as that of the first treatment. Together, these results indicate that PFQX1(AF488) can be used to visualize endogenous AMPARs in cultured hippocampal neurons in a rapid, reversible, and reproducible manner.

### Fluorescent visualization of cell-surface AMPARs during synaptic plasticity

Many studies have reported that neuronal AMPAR trafficking is dynamically regulated during synaptic activity^7–9^. In particular, AMPAR accumulation at the postsynaptic terminal has been observed with LTP stimulation. However, the current techniques largely depend on the fusion of fluorescent proteins to AMPARs, and in most cases, the fusion proteins are overexpressed in neurons. Ideally, endogenous AMPARs should be visualized under live conditions with minimal perturbation. Taking advantage of the unique binding features of PFQX1(AF488), we thus used PFQX1(AF488) to investigate endogenous AMPAR trafficking in neuronal cells both before and after LTP stimulation.

In the hippocampus, LTP is initiated by the activation of *N*-methyl-D-aspartate-type glutamate receptors (NMDARs) in response to repetitive neuronal stimulation^1,32,33^. The selective activation of NMDARs using chemical compounds can also induce LTP^34,35^, which is known as chemically induced LTP (cLTP). Here, we used PFQX1(AF488) to investigate cell-surface AMPAR levels before and after cLTP stimulation. As shown in Fig. 5a, cell-surface AMPARs were stained with PFQX1(AF488), and fluorescent images were obtained as the basal state using confocal microscopy. Following the removal of PFQX1(AF488) by medium exchange, cLTP stimulation was applied. To induce cLTP, Mg^2+^ in the medium was withdrawn in conjunction with the addition of 200 μM glycine, a co-agonist of NMDAR, for 15 min for the selective activation of NMDAR. After medium exchange to the incubation buffer and 1 hour of incubation, PFQX1(AF488) was again added to stain cell-surface AMPARs; fluorescent images after cLTP stimulation were then obtained using confocal microscopy (**Fig. 5b**). The fluorescent intensity of PFQX1(AF488) in spines was prominently enhanced after cLTP stimulation, and the quantification of PFQX1(AF488) fluorescent intensity in spines showed a 1.6-fold increase compared with the basal state (**Fig. 6c**). By contrast, the fluorescent intensity of PFQX1(AF488) did not significantly change after 1 hour of incubation in cells without cLTP stimulation. Recent studies have suggested that remodeling of the synaptic microenvironment occurs during LTP, which may affect the ligand-binding properties of AMPARs or alter the pH of synaptic regions^26^. In this context, the affinity of PFQX1(AF488) to AMPARs was unchanged even after cLTP stimulation (**Extended Data Fig. 8a**). In addition, as described in the first Results subsection, the fluorescent intensity of PFQX1(AF488) was almost unaffected at pH 5–10 (**Extended Data Fig. 4**). These data suggest that the enhancement of fluorescent intensity after cLTP stimulation would be attributed to an increase in the number of AMPARs at the synaptic terminal.

**Fig. 5.**
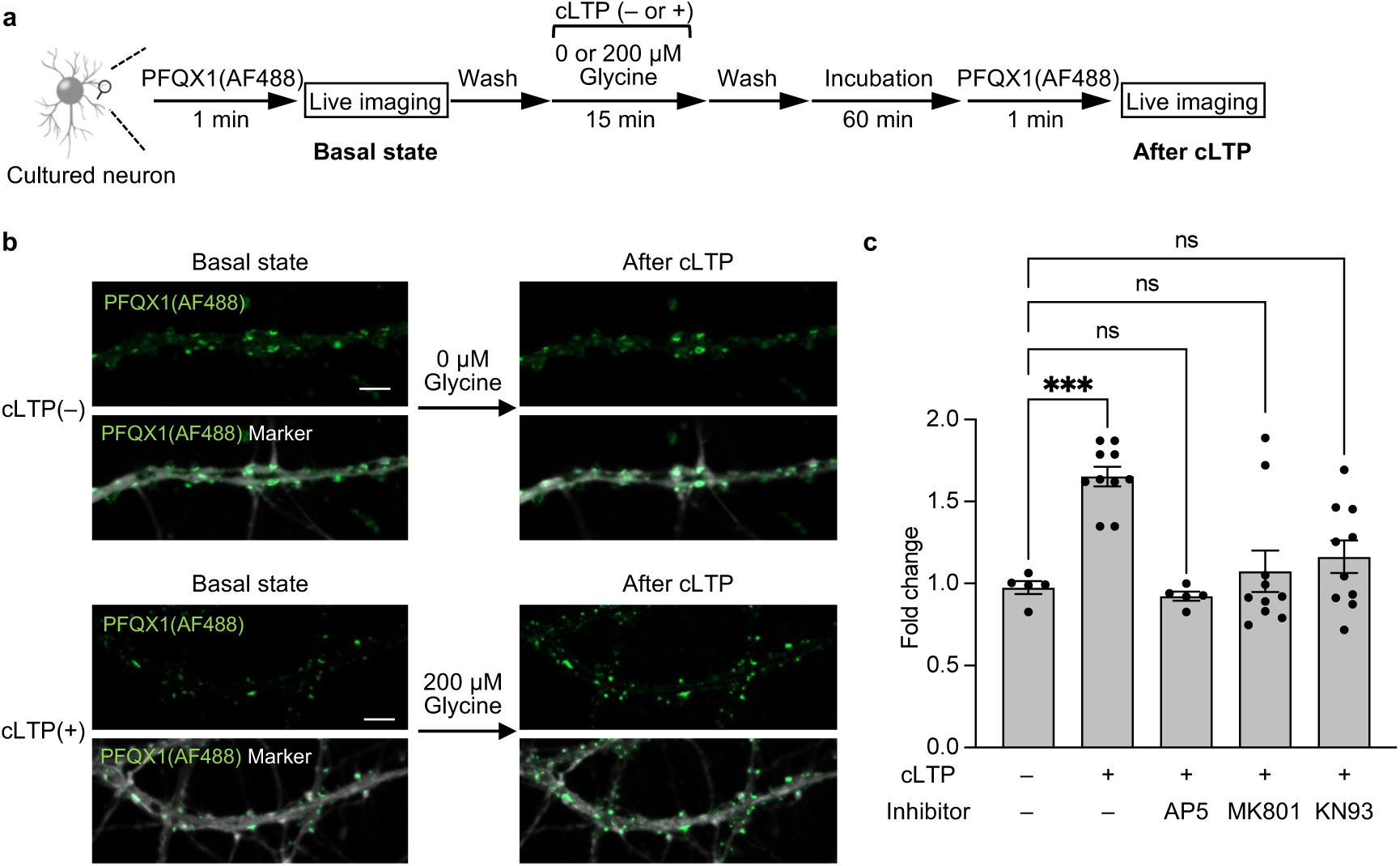
Visualization of the surface accumulation of AMPARs by cLTP stimulation. **(a)** Experimental scheme of cLTP stimulation. “Basal state” image were obtained after the addition of 100 nM PFQX1(AF488) to cultured hippocampal neurons. The probe was washed out by medium exchange. Then, cLTP was induced in cLTP stimulation buffer containing 200 µM glycine at room temperature for 15 min. After medium exchange with cLTP incubation buffer, the cells were further incubated for 60 min. Then, the cells were treated again with 100 nM PFQX1(AF488) to obtain “after cLTP” images. **(b)** Representative images of the cLTP experiment. Upper, images of the control condition “cLTP(–)” where glycine was not added to the cells are shown. Lower, images of the cells stimulated with glycine “cLTP(+)” are shown. Calcein red signals are shown in gray as a cytoplasmic marker. Scale bars, 5 µm. **(c)** The intensity of PFQX1(AF488) in spines was analyzed by the intensity of “after cLTP” images divided by that of “basal state” images. [AP5] = 50 µM, [MK-801] = 20 µM and [KN93] = 5 µM (n = 5 – 10). ***Significant difference from cLTP(–) (p < 0.001, One-way ANOVA with Dunnet’s test). ns, not significant (p > 0.05). Data are represented as mean ± s.e.m.

Both NMDAR activation and the subsequent phosphorylation of synaptic proteins by Ca^2+^/calmodulin-dependent protein kinase II (CaMKII) are critical steps in the expression of LTP^1,32,33,36,37^. We therefore examined whether the increase in PFQX1(AF488) fluorescence after cLTP stimulation was indeed correlated to the activation of both NMDARs and CaMKII. As shown in Fig. 5c, pretreatment with NMDAR blockers (AP5 or MK-801) or a CaMKII inhibitor (KN93) failed to increase PFQX1(AF488) fluorescence, even after cLTP stimulation. This result indicates that the observed increase in PFQX1(AF488) fluorescence is strongly correlated with Ca^2+^ influx through NMDARs and the resulting downstream pathway.

One possible mechanism for the observed increase in the number of AMPARs on synaptic membranes is an elevation in AMPAR expression. We therefore quantified AMPAR expression levels before and after cLTP stimulation using western blotting. As shown in **Extended Data Fig. 8b**, there were no prominent changes in the expression levels of GluA1 or GluA2 (the main subunits of AMPARs in the hippocampus) even after cLTP stimulation^38^. Thus, the increased fluorescent intensity of PFQX1(AF488) in spines can likely be attributed to altered AMPAR trafficking in neurons.

### Quantitative analysis of AMPAR trafficking during synaptic plasticity

For the delivery of AMPARs to synaptic membranes, two possible pathways have been proposed: the insertion of proteins from intracellular spaces by exocytosis^39,40^, and the lateral diffusion of proteins from the surface of dendritic regions^11,41,42^ (**Fig. 6a**). However, the contribution of both pathways during LTP remains controversial (see the Discussion for details). In the present study, the use of PFQX1(AF488) revealed increased AMPARs in spines after cLTP stimulation by allowing the visualization of cell-surface AMPARs before and after cLTP stimulation (**Fig. 5**). Although this technique is powerful for visualizing cell-surface AMPARs in each period, it does not give any information about the trafficking pathway. We have previously reported a method for the covalent chemical labeling of cell-surface AMPARs, termed ligand-directed two-step labeling (**Extended Data Fig. 1b**). However, it is unable to provide information regarding intracellular AMPARs, while this covalent labeling method is useful for tracking labeled AMPARs on plasma membranes. Taking advantage of the key features of each method, we therefore combined PFQX1(AF488) with the two-step labeling method to elucidate the trafficking mechanism of AMPARs during cLTP.

**Fig. 6.**
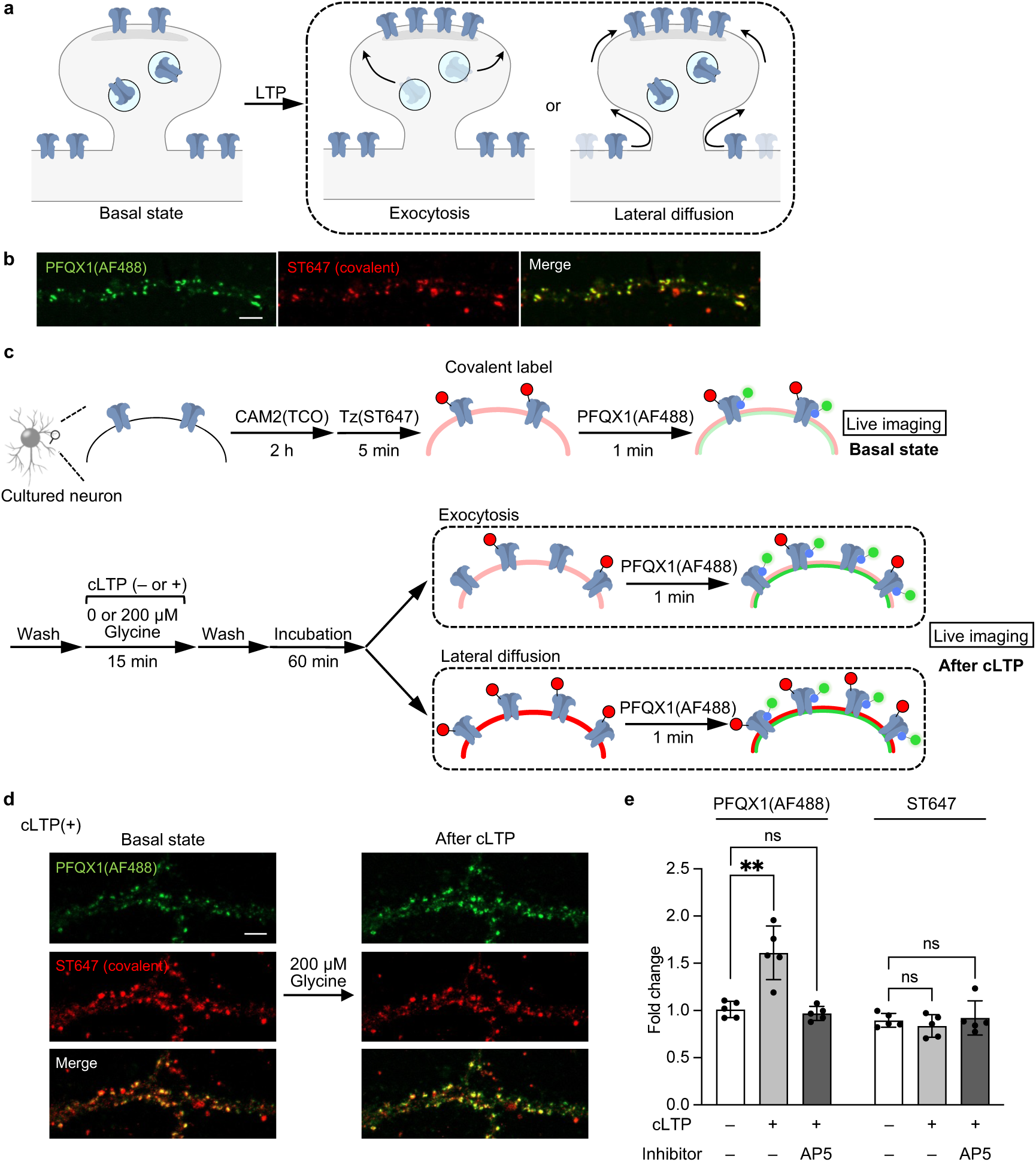
Mechanisms of the surface accumulation of AMPARs by cLTP stimulation. **(a)** Putative mechanisms of AMPAR accumulation in synapses during LTP. **(b)** Confocal live imaging of cultured hippocampal neurons. The cells were treated with 2 µM CAM2(TCO) for 2 h, followed by the addition of 100 nM Tz(ST647) for 5 min. Then, 100 nM PFQX1(AF488) was added to the cells. A detailed scheme of two-step labeling of ST647 is shown in Extended Data Fig. 9a. Scale bar, 5 µm. **(c)** A schematic illustration of the experimental procedure for dicreminating between exocytosis or lateral diffusuion is shown. Cultured hippocampal neurons were covalently labeled with 2 µM CAM2(TCO) for 2 h followed by 100 nM Tz(ST647) for 5 min. “Basal state” images were scanned after the addition of 100 nM PFQX1(AF488). After washing out of PFQX1(AF488) by medium exchange, cLTP was induced by 200 µM glycine for 15 min, and the cells were further incubated for 60 min in cLTP incubation buffer. The cells were treated with 100 nM PFQX1(AF488) again to obtain “after cLTP” images. If exocytosis was the main pathway of AMPAR accumulation, the covalent labeling signal by ST647 would not be changed. If lateral diffusion was the main pathway, both ST647 and PFQX1(AF488) signals would be enhanced. A schematic illustration of putative results regarding ST647 labeling is also shown in Extended Data Fig. 9c. **(d)** Representative live cell images of the cLTP experiment. Images of the cells stimulated with glycine are shown. Scale bar, 5 µm. **(e)** The intensity of AF488 and ST647 in spines was analyzed. Fluorescent intensity of “after cLTP” images was divided by that of “basal state” images. Raw images for cLTP(–) or pretremement of AP5 is shown in Extended Data Fig. 9d or 9e, respectively. [AP5] = 50 µM (n = 5). **Significant difference (p < 0.01, One-way ANOVA with Dunnet’s test). ns, not significant (p > 0.05). Data are represented as mean ± s.e.m.

Prior to analyzing AMPAR trafficking, we first examined the feasibility of using PFQX1(AF488) in conjunction with the two-step labeling method. To demonstrate dual staining, cell-surface AMPARs were covalently labeled with a fluorophore, SeTau-647 (ST647), by two-step labeling; specifically, cultured hippocampal neurons were labeled with CAM2(TCO) followed by treatment with 100 nM Tz(ST647) for 5 min (**Extended Data Fig. 9a**). After the two-step labeling, the same neurons were stained with 100 nM PFQX1(AF488). As shown in Fig. 6b, the fluorescent signal of PFQX1(AF488) merged well with that of ST647. In addition, immunostaining of the neurons after fixation revealed that the ST647 signals were observed alongside the dendrite marker microtubule-associated protein 2 (MAP2) (**Extended Data Fig. 9b**), which merged well with the postsynaptic marker PSD95. Together, these findings suggest that AMPARs are selectively labeled with ST647, and indicate that the two-step labeling method with PFQX1(AF488) staining is successfully able to simultaneously visualize AMPARs in living hippocampal neurons.

We then analyzed changes in AMPAR trafficking after cLTP stimulation by combining the two-step labeling and PFQX1(AF488). The experimental procedure and two possible results are shown in Fig. 6c. If exocytosis was the main mechanism of synaptic accumulation, unlabeled AMPARs would be increased at the synaptic surface, thus resulting in increased PFQX1(AF488) signals but no changes in ST647 intensity (**Fig. 6c** and **Extended Data Fig. 9c**). Conversely, if lateral diffusion was the main mechanism, labeled AMPARs would enter synapses from extrasynaptic regions, thus resulting in increases in both ST647 and PFQX1(AF488) fluorescent intensity in the spines. As shown in Fig. 6d and 6e, cLTP stimulation increased the fluorescence of PFQX1(AF488) but failed to increase ST647 fluorescence in spines. Importantly, in the absence of cLTP stimulation, neither PFQX1(AF488) nor ST647 showed significant changes in fluorescent intensity (**Fig. 6e** and **Extended Data Fig. 9d**). Even after cLTP stimulation, pretreatment with the NMDAR inhibitor AP5 failed to increase both PFQX1(AF488) and ST647 fluorescence (**Fig. 6e** and **Extended Data Fig. 9e**). Moreover, as shown in **Extended Data Fig. 10**, no significant fluorescence changes were observed even if AMPARs were covalently labeled with AF488 using two-step labeling with Tz(Ax488) instead of Tz(ST647). This indicated that the differences in fluorescence changes after cLTP stimulation between PFQX1(AF488) and ST647 labeling were unlikely to be caused by differences in fluorophores. Overall, our results strongly indicate that exocytosis is the main mechanism of AMPAR accumulation in dendritic spines during cLTP in cultured hippocampal neurons.

## Discussion

In the present study, we developed PFQX1(AF488) for the rapid and reversible staining of AMPARs, which allowed us to quantify cell-surface AMPARs in living cells. Using PFQX1(AF488), AMPAR accumulation in spines during synaptic plasticity was successfully visualized by obtaining images both before and after cLTP stimulation in cultured hippocampal neurons. Furthermore, the trafficking mechanisms of AMPARs during synaptic plasticity were quantitatively evaluated using both PFQX1(AF488) and the chemical labeling of ST647 to cell-surface AMPARs. This analysis revealed that cLTP stimulation-induced AMPAR accumulation in spines was mainly mediated by exocytosis from the intracellular compartment rather than by lateral diffusion on the cell surface.

The affinity of PFQX1(Dye) to AMPARs was highly dependent on the structure of the conjugated fluorophore; both PFQX1(Fl) and PFQX1(AF488) had high affinity, whereas PFQX1(TAMRA) had low affinity. These differences were revealed in the current study via an X-ray crystallographic analysis of the LBD of AMPAR complexed with PFQX1(AF488). In the X-ray structure, AF488 formed hydrophobic and electrostatic interactions with the LBD. Consistent with the results of the structural analysis, the introduction of a linker between the fluorophore and the PFQX ligand in PFQX2(Fl) resulted in a marked decrease in AMPAR affinity. This structural information provides important insights for the molecular design of new fluorescent probes to AMPARs, such as probes with long-wavelength-emitting dyes that are suitable for visualizing endogenous AMPARs in brain tissue or in vivo.

Staining with PFQX1(AF488) was a very simple procedure, and AMPARs were able to be visualized immediately after the addition of probes to the culture medium. Moreover, given that PFQX1(AF488) is an antagonist-based probe, the function of stained AMPARs was inhibited during visualization. However, because of its relatively moderate affinity (*K_d_* = 37.2 nM), PFQX1(AF488) was able to be readily removed by a simple medium exchange procedure. Moreover, AMPARs were visualized with high reproducibility by re-treatment with the same concentration of PFQX1(AF488). Together, these features allowed us to quantify the changes in AMPAR trafficking during cLTP with minimal perturbation of AMPARs. This method will therefore be powerful for obtaining snapshot images of AMPAR dynamics under live conditions without any genetic manipulation.

In the present study, PFQX1(AF488), bearing AF488 dye, was selected as a probe for visualizing AMPARs in hippocampal neurons because it allowed us to visualize the receptors in a pH-independent manner in the pH 5–10 range. Another probe, PFQX1(Fl), bearing F1, also showed high affinity to AMPARs. However, in contrast to PFQX1(AF488), the fluorescent intensity of PFQX1(Fl) was weakened under acidic conditions (pH < 6). This property will be advantageous for evaluating AMPAR dynamics during long-term depression, which is another form of synaptic plasticity. Long-term depression refers to a long-lasting decrease in the efficacy and strength of synaptic transmission. Several mechanistically distinct forms of this type of synaptic plasticity have been identified, and the key step is believed to be the endocytosis of AMPARs into acidic organelles, such as endosomes and lysosomes^5,15^. Thus, an AMPAR probe with pH sensitivity will likely be useful for analyzing the internalization of endogenous AMPARs.

To date, two mechanisms have been proposed for the accumulation of AMPARs to the postsynaptic surface during LTP. The first is exocytosis from intracellular endosomes at extrasynaptic, peri-synaptic, or synaptic regions^39,40^. The other is long-range lateral diffusion on the cell surface from the dendrites or extrasynaptic regions^11,41,42^. However, it has remained controversial whether exocytosis or lateral diffusion is the main mechanism, mainly because of a lack of appropriate methods for evaluating the trafficking of AMPARs endogenously expressed in neurons without perturbing the receptors or neuronal function. To visualize AMPAR trafficking, SEP-tagged AMPARs on the N-terminus (SEP-AMPARs) have been frequently used. SEP fluorescence is weakened under acidic conditions, which allows for the visualization of exocytosis steps from acidic endosomes. However, although SEP-AMPARs are potent, they are overexpressed in most cases, which raises concerns that AMPAR distribution is affected. In addition, post-translational modifications or associations with accessory proteins may be affected. It has also been suggested that fluorescent changes of SEP during LTP is caused by changes in intracellular pH rather than AMPAR localization changes^43^. In contrast to previous techniques, our method was able to visualize the trafficking changes of endogenous AMPARs during LTP with minimal perturbation. We therefore believe that the results of the current study, in which exocytosis was identified as the main pathway, are highly reliable. Given its relatively simple procedures, this technique is expected to be widely applied to other neuronal cultures, including brain slices, to clarify the relationship between AMPAR trafficking changes and neuronal functions.

## Methods

### Synthesis

All synthesis procedures and compound characterizations are described in Supplementary information.

### Construction of expression plasmids

cDNAs of mouse GluA1^flip^(Q), 5’UTR - GluA1^flip^(Q) - 3’UTR, GluA2 ^flip^(Q), GluA2^flip^(Q) - 3’UTR, GluA3 ^flip^(Q), and GluA4 ^flip^(Q) were amplified from mouse brain marathon-ready cDNA (Clontech) by PCR. These cDNAs were subcloned into the expression vector, pCAGGS (kindly provided by Dr J. Miyazaki, Osaka University, Osaka, Japan) or pCDM. The pCDM vector was prepared from pcDNA3.1(+) (Invitrogen), in which the neomycin cassette was excised using *Pvu*II. 5’UTR - GluA1^flip^(R) - 3’UTR and GluA3 ^flip^(Q) (Y454A/R461G) was obtained using the Q5® Site-Directed Mutagenesis Kit (NEB). To obtain Halo-tag - GluA2^flip^(Q) - 3’UTR (Halo-GluA2), cDNA encoding Halo-tag was amplified from pFN21A Halo-tag® CMV Flexi® Vector (Promega). The Halo-tag cDNA was inserted into the pCAGGS expression vector before GluA2^flip^(Q) - 3’UTR.

The sequence of genetically engineered ligand binding domain named S1S2J was designed as the residues S383-K506 and P632-S775 of matured GluA2 conjugated through Gly and Thr as a linker, as previously reported^28^. Besides, the three mutations G389R, L390G, and E391A were included. This gene fragment encoding the S1S2J was cloned into the modified T7 expression vector^44^. The expression construct contained the additional Met and Gly to its N-terminus, while the hexa-histidine tag was not included at the N-terminus. All PCR-amplified DNAs were confirmed by DNA sequence analyses.

### Cell culture and transfection

HEK293T cells (ATCC) were cultured in Dulbecco’s modified Eagle’s medium (DMEM)-GlutaMAX (Invitrogen) supplemented with 10% FBS (Sigma-Aldrich), penicillin (100 units ml^−1^), and streptomycin (100 µg ml^−1^), and incubated in a 5% CO_2_ humidified chamber at 37 °C. Cells were transfected with a plasmid using Lipofectamine 3000 (Invitrogen) according to the manufacturer’s instructions. After 24 h from transfection, the cells were seeded on glass bottom dishes coated with poly-L-lysine solution and incubated for 24 h at 37°C in a humidified atmosphere of 95% air and 5% CO_2_.

### Confocal live cell imaging of cell-surface AMPARs in HEK293T cells

HEK293T cells were co-transfected with AMPAR plasmid (GluA1, GluA2, or the control vector) and transfection marker (EFGP-F of mCherry-F). Fluorophore–ligand conjugates were diluted in HBS buffer (20 mM HEPES pH 7.4, 107 mM NaCl, 6 mM KCl, 1.2 mM MgSO_4_, 2 mM CaCl_2_, 11.5 mM glucose), and added to the cells. Confocal live imaging was performed with LSM900 (Carl Zeiss) equipped with a 63×, numerical aperture (NA) = 1.4 oil-immersion objective. Fluorescence images were acquired by excitation at 405, 488, 561, or 640 nm derived from diode lasers.

For competitive inhibition assay by NBQX, the stained cells were scanned first, and then 10 µM NBQX was added to the dish to inhibit probe binding.

For evaluating the concentration-dependency of fluorophore–ligand conjugates, 0.1 nM probe in HBS was added to the dish. Then, probes in HBS were step-wisely added to the dish to a final concentration of 0.3, 1, 3, 10, 30, 100, 300, 1000 and 3000 nM. Cell surface fluorescence intensity was quantified from the line scans of transfection marker positive cells and calculated with subtraction of background by ZEN blue software (Carl Zeiss). The membrane intensity was fitted with KaleidaGraph (Synergy software) to calculate the *K_d_* value using the following equation: a + (b-a)/(1+(x/c)^d).

### Evaluation of the affinity of PFQX-amine

HEK293T cells were co-transfected with GluA2 and mCherry-F as a transfection marker. After washing the cells with HBS, 30 nM PFQX1(AF488) was added to the dish. Then, PFQX-amine in HBS was step-wisely added to the dish to a final concentration of 1, 3, 10, 30, 100, 300, 1000, 3000, 10000 and 30000 nM. Imaging was performed with confocal microscopy. Cell surface fluorescence intensity was quantified from the line scans of transfection marker positive cells calculated by ZEN blue software and calculated with subtraction of background. The membrane intensity was fitted with KaleidaGraph to calculate the *IC_50_* value using the following equation: a + (b-a)/(1+(x/c)^d). The *K_i_* value was calculated using the Cheng-Prusoff equation: *K_i_* = *IC_50_*/(1+[S]/*K_d_*). In the equation, [S] and *K_d_* were the concentration and dissociation constant of PFQX1(AF488), respectively.

### pH-dependency of fluorescence intensity

Fluorescence intensity was determined by spectrofluorophotometer RF-6000 (Shimadzu). PFQX1(Fl) or PFQX1(AF488) was added to the 0.1 M GTA buffer (0.1 M 3,3-dimethyl glutaric acid, 0.1 M Tris, 0.1 M 2-amino-2-methyl-1,3-propanediol) to a final concentration of 2 µM. pH value of GTA buffer was adjusted to 4, 5, 6, 7, 8, 9, and 10. Fluorescence intensity was normalized to the intensity at pH 10.

### Expression and purification of S1S2J

The expression plasmids were transferred into *E. coli* BL21(DE3) expression host cells. The cells were initially grown in 5 mL LB medium supplemented with 50 μg/mL kanamycin at 37 °C for a day. 800 μL of the cultured cells were then transferred to 800 mL LB medium supplemented with 50 μg/mL kanamycin. The cells were cultured at 37 °C until the O.D._600_ reached 0.6. At this point, 1 mM IPTG (isopropyl-β-_D_-thiogalactopyranoside) was added to the medium for expression of S1S2J, and the cells were cultured at 37 °C for an additional 2 hours. The cells were harvested by centrifugation and stored at −80 °C until use.

The cells were resuspended to 10 mL of buffer A (50 mM Tris-HCl pH 8.0, 100 mM NaCl, 1 mM EDTA, 1 mM PMSF, 3 mM sodium deoxycholate, 1 mg/mL lysozyme, 40 μg/mL DNase I) and disrupted by sonication. After 15 min incubation at 4 °C, the suspension was centrifuged at 17,000 × *g*, and the insoluble fraction was collected as the crude inclusion bodies. As followed by the same centrifugal separation, the crude inclusion bodies were washed with 20 mL of buffer B (50 mM Tris-HCl pH 8.0, 100 mM NaCl, 10 mM EDTA, 1 mM PMSF, 0.5% (v/v) Triton X-100), and then buffer C (20 mM Tris-HCl pH 7.4, 200 mM NaCl, 1 mM EDTA, 1 mM PMSF). The prepared inclusion bodies were stored at −80 °C until use.

The inclusion bodies were solubilized to 15 mL of buffer D (50 mM Tris-HCl pH 7.4, 5 mM EDTA, 8 M guanidine hydrochloride, 50 mM DTT) by stirring at room temperature for 4 hours. Then, 100 mL of cold buffer E (20 mM sodium acetate, pH 4.5, 1 mM EDTA, 4 M guanidine hydrochloride, 1 mM DTT) was added to the denatured proteins and stirred for 1 hour at 4 °C. The supernatant was collected after centrifugation at 17,000 × *g* for 1 hour, and the protein concentration was adjusted to 1.0–1.5 mg/mL with buffer E. This solution was dialyzed against 2 L of buffer F (10 mM NaCl, 0.4 mM KCl, 650 mM arginine hydrochloride, 1 mM EDTA, pH 8.5) at 4 °C for 16 hours (2 times). The solubilized proteins were centrifuged at 17,000 × *g* for 1 hour, and the supernatant was collected. Then, 1 mM reduced glutathione and 0.2 mM oxidized glutathione were added to the supernatant and incubated at 4 °C for a day. The solution was centrifuged at 17,000 × *g* for 1 hour, and the supernatant was collected. The supernatant was dialyzed against 2 L of three different buffers in the following order: the buffer containing buffer F and buffer G (10 mM HEPES-NaOH pH 7.4, 200 mM NaCl, 1 mM EDTA) at a volume ratio of 1:1 at 4 °C for 4 hours, the buffer containing buffer F and buffer G at a volume ratio of 1:9 at 4 °C for 4 hours, and buffer G at 4 °C for a day. After dialysis, the supernatant was separated from the precipitant by centrifuges at 17,000 × *g* for 1 hour. The supernatant was concentrated using Centricon Plus-70 with a 30 kDa cutoff (Millipore). The concentrated fraction was loaded onto HiLoad 26/600 superdex 75 column (Cytiva) equilibrated with buffer G. After collecting the eluate containing the S1S2J, the buffer was exchanged to buffer H (10 mM HEPES-NaOH pH 7.0, 20 mM NaCl, 1 mM EDTA) by repetitive centrifugal concentration and dilution using Amicon Ultra 4 with 30 kDa cutoff and concentrated to 10 mg/mL.

### Crystallization and X-ray diffraction data collection

To prepare the S1S2J bound to the ligand PFQX1(AF488), the purified S1S2J was mixed with 1 mM of PFQX1(AF488) at 4 °C for a day. The crystallization was performed by sitting-drop vapor-diffusion method using the JCSG-plus Eco screen (Molecular Dimensions). A crystallization robot, Gryphon (Art Robins Instruments), was used to mix 0.3 μL of the purified S1S2J with an equal volume of reservoir solution. The single crystal of S1S2J coloured in yellow grew to dimensions of 0.7 × 0.2 × 0.2 mm in the reservoir (0.2 M lithium sulfate, 0.1 M Tris-HCl, pH 8.5, 40% (v/v) PEG400) within a day at 4 °C. The crystal was frozen with liquid nitrogen. Diffraction data were collected on beamline BL45XU at SPring-8 (Hyogo, Japan) and automatically processed using the *ZOO* system^45^. Data statistics are summarized in **Supplementary Table 1**.

### Structure determination and refinement

The structure of S1S2J bound to PFQX1(AF488) was determined by the molecular replacement (MR) method with *AutoMR* from the *Phenix* program package^46^. The structure of the S1S2J in complex with antagonist 2,3-dioxo-6-nitro-1,2,3,4-tetrahydrobenzoquinoxaline-7-sulfonamide (NBQX) (PDB code: 6FQH) was used as the search model. Rotation and translation functions were calculated using the data of 47.3-2.5 Å resolution. Subsequent cycles of refinement were conducted using *phenix.refine* from the *Phenix* suite, alternating with manual fitting and rebuilding of the macromolecules, ions, polymers and waters based on 2*F*_O_−*F*_C_ and *F*_O_−*F*_C_ electron density maps using *Coot*^47^. Then, PFQX1(AF488) was built based on *F*_O_−*F*_C_ electron densities, and validated based on the polder omit map^48^. The final refinement statistics and geometry, as defined by *MolProbity*^49^, are presented in Supplementary Table 1. The structural figures were generated using *PyMOL* (http://www.pymol.org).

### Kinetics and reversibility of PFQX1(AF488) binding in HEK293T cells

HEK293T cells were transfected with Halo-tag-fused GluA2. The cells were incubated with 500 nM Halo-Tag ligand-Alexa647 (HTL (AF647)) for 15 min at room temperature. AF647 was utilized as a transfection marker. The cells were washed with HBS three times, and a cloning ring (16 mm φ) was placed on the dish to accelerate the diffusion of the probe solution. After the dish was set on the stage, 100 nM PFQX1(AF488) was added to inside the cloning ring and imaged at specified time points by confocal microscopy.

For evaluating reversibility, 100 nM PFQX1(AF488) in HBS was added to the dish. The cells were observed by confocal microscopy as the first image. Then, the cells were washed with HBS three times and scanned as the wash image. After removal of the buffer, 100 nM PFQX1(AF488) was re-treated, and the cells were scanned as the second image. To quantify the fluorescence intensity of the membrane, the average signal intensity of ROIs set on the membrane was calculated after subtracting background fluorescence by ZEN blue software.

### Fluorescence Ca^2+^ imaging

HEK293T cells transfected with Ca^2+^-permeable GluA1–4 (GluA1^flip^(Q), GluA2^flip^(Q), GluA3^flip^(Q) (Y454A/R461G) and GluA4^flip^(Q)) were seeded on the 96 well plate. The cells were loaded with 5 µM Cal-520 AM (AAT Bioquest) for 2 h in the culture medium and washed with HBS using AquaMax 2000 (Molecular devices). The imaging experiment was carried out in an HBS buffer. For the fluorescence Ca^2+^ imaging, PFQX-amine was pretreated to the cells for 3 min followed by the addition of 100 µM CTZ with glutamate to inhibit the desensitization. Fluorescence intensity of Cal-520 was obtained using FlexStation3 (Molecular devices), and analyzed with SoftMax Pro 7 (Molecular devices). ΔRFU was defined as the difference between the maximum intensity value after adding glutamate and the average intensity value before adding the glutamate. ΔRFU was fitted with KaleidaGraph to calculate *IC_50_* value using the following equation: a + (b-a)/(1+(x/c)^d).

### Animals

Pregnant ICR mice maintained under specific pathogen-free conditions were purchased from Japan SLC, Inc (Shizuoka, Japan). The animals were housed in a controlled environment (23 ± 1 °C, 12 h light/dark cycle) and had free access to food and water, according to the regulations of the Guidance for Proper Conduct of Animal Experiments by the Ministry of Education, Culture, Sports, Science, and Technology of Japan. All experimental procedures were performed in accordance with the National Institute of Health Guide for the Care and Use of Laboratory Animals, and were approved by the Institutional Animal Use Committees of Nagoya University.

### Preparation of primary cultured hippocampal neurons

We modified the banker’s culture method of hippocampal neurons^50^. Glass bottom dishes (14 mm φ, Matsunami) were coated with poly-D-lysine (Sigma-Aldrich) and washed with sterile dH_2_O three times. Hippocampi from 16-day-old ICR mouse embryos were aseptically dissected and digested with 0.25% (w/v) trypsin (Nacalai Tesque) for 20 min at 37°C. The cells were resuspended in Neurobasal Plus medium supplemented with 10% FBS, 1.0 mM GlutaMAX I (Invitrogen), penicillin (100 units/ml), and streptomycin (100 µg/ml) and filtered by Cell Strainer (100 µm, Falcon) followed by centrifugation at 1,000 rpm for 5 min. The cells were resuspended in Neurobasal Plus medium supplemented with 2% of B-27 Plus Supplement, 1.0 mM GlutaMAX I (Invitrogen), penicillin (100 units/ml), and streptomycin (100 µg/ml) and plated at a density of 2 × 10^4^ cells on glass bottom dishes. Precultured cortical astrocytes on glass coverslips (13 mm φ) were transferred to the dishes so that astrocytes were faced to neurons. The cultures were maintained at 37 °C in a 95% air and 5% CO_2_ humidified incubator. The culture medium was replaced every 7 days, and the neurons were used at 16–18 DIV.

### Confocal live cell imaging of AMPARs in cultured neurons

50 nM Calcein Red AM (AAT Bioquest) was added to the cultured neurons for 30 min at 37 °C to visualize the intracellular region. The cells were washed with HEPES-based ACSF (25 mM HEPES pH 7.4, 140 mM NaCl, 5 mM KCl, 1.3 mM CaCl_2_, 2 mM MgCl_2_, 33 mM glucose), and 100 nM PFQX1(AF488) in HEPES-based ACSF was added to the dish.

For evaluating the concentration-dependency of PFQX1(AF488), the cells were treated with 1 nM PFQX1(AF488). Then, PFQX1(AF488) in HEPES-based ACSF was step-wisely added to the dish to a final concentration of 3, 10, 30, 100, 300 and 1000 nM. To quantify the concentration-dependency after cLTP, the procedure described above was conducted after cLTP stimulation as described in “cLTP and quantification of cell-surface AMPARs”. The average intensity of ROIs set on spines was calculated by ZEN blue software, after subtracting background fluorescence. The intensity was fitted with KaleidaGraph to calculate the *K_d_* value using the following equation: a + (b-a)/(1+(x/c)^d).

### Kinetics and reversibility of PFQX1(AF488) binding in cultured neurons

A cloning ring (16 mm φ) was placed on the dish to accelerate the diffusion of the probe solution. After the dish was set on the stage, the cells were treated with 100 nM PFQX1(AF488) and imaged at specified time points by confocal microscopy.

For evaluating reversibility, stained cells were observed by confocal microscopy as the first image. Then, the cells were washed with HEPES-based ACSF three times and scanned as the wash image. After removal of the buffer, 100 nM PFQX1(AF488) was re-treated, and the cells were scanned as the second image. To quantify the fluorescence intensity of spines, the average signal intensity of ROIs set on spines was calculated by ZEN blue software after subtracting background fluorescence.

### Immunostaining of cultured neurons

The cultured hippocampal neurons were fixed with cold methanol for 5 min and washed with PBS three times. Fixed cells were incubated with Blocking One Histo (Nacalai Tesque) for 10 min. The cells were washed with PBS containing 0.05% Tween 20 (PBS-T) once and incubated with primary antibodies in PBS-T containing 5% Blocking One Histo at room temperature for 2 h. Then, the cells were washed with PBS-T three times and incubated with secondary antibodies in PBS-T containing 5% Blocking One Histo for 1 h at room temperature. Following primary antibodies were used: mouse anti-PSD95 (abcam, ab2723, 1:1,000) or rabbit anti-MAP2 (Millipore, AB5622, 1:1,000). The following secondary antibodies were used: donkey anti-mouse CF568 conjugated (Sigma, SAB4600075-250UL, 1:1,000), goat anti-mouse Alexa Fluor 647 conjugated (Invitrogen, A21235, 1:1,000), goat anti-rabbit Alexa Fluor 405 conjugated (abcam, ab175652, 1:1,000) and anti-rabbit Alexa Fluor Plus 488 conjugated (Invitrogen, A32731, 1,000). Imaging was performed with confocal microscopy.

With two-step labeling, the cells were labeled as described in “Two-step labeling and PFQX1(AF488) staining in cultured neurons,” followed by immunostaining.

In Extended Data Fig. 7a, the cells were incubated with 500 nM CellTracker Red CMPTX (Invitrogen) for 30 min at 37 °C to visualize the intracellular region before the immunostaining.

### cLTP and quantification of cell-surface AMPARs

After washing the cells with HEPES-based ACSF, 100 nM PFQX1(AF488) was added to the dish. The cells were scanned to obtain “basal state” images. Then, the cells were washed with cLTP stimulation buffer (HEPES-based ACSF except for MgCl_2_ supplemented with 20 µM bicuculline and 1 µM strychnine) three times to remove PFQX1(AF488) cLTP was induced by 200 µM glycine in cLTP stimulation buffer for 15 min at room temperature. After washing the cells with cLTP incubation buffer (HEPES-based ACSF supplemented with 20 µM bicuculline and 1 µM strychnine), the cells were incubated for 60 min at room temperature. Finally, 100 nM PFQX1(AF488) was added to the dish, and images were scanned to obtain “after cLTP” images. Inhibitors (50 µM AP5, 20 µM MK-801 for NMDARs and 5 µM KN93 for CaMKII) were added to cLTP stimulation buffer and cLTP incubation buffer. To quantify the fluorescence intensity in spines, the average signal intensity of ROIs set on spines was calculated by ZEN blue software after subtracting background fluorescence. The fold change value was calculated using the following equation: (fluorescence intensity of “after cLTP” images) / (fluorescence intensity of “basal state” images).

### Western blotting for quantification of AMPARs

cLTP was performed as described in “cLTP and quantification of cell-surface AMPARs”. After washed with HBS, the cells were lysed with 100 µL of radio immunoprecipitation assay (RIPA) buffer (50 mM Tris-HCl pH 7.4, 150 mM NaCl, 1% NP-40, 0.5% sodium deoxycholate, 0.1% SDS, and 1 mM EDTA) containing 1% protease inhibitor cocktail (Nacalai Tesque) and incubated for 30 min at 4°C on a mini rotary incubator. After mixing with 5 × Laemmli sample buffer (300 mM Tris-HCl pH 6.8, 15% SDS, 19.7% sucrose, and 0.05% bromophenol blue) containing 250 mM DTT, the samples were incubated for 30 min at room temperature with shaking. The samples were applied to SDS-PAGE (BIO-RAD Mini-Protean III) and electrotransferred onto an immuno-blot polyvinylidene fluoride membrane, followed by blocking with 5% nonfat dry milk in Tris-buffered saline (10 mM Tris-HCl pH 8.0, 150 mM NaCl) containing 0.05% Tween 20 (TBS-T). The membrane is incubated with primary antibodies in TBS-T supplemented with 1% nonfat dry milk overnight at 4 °C. Samples were detected by rabbit anti-GluA1 antibody (abcam, ab19491, 1:3,000), rabbit anti-GluA2 antibody (abcam, ab206293, 1:3,000), and rabbit βIII tubulin antibody (abcam, ab18207, 1:3,000). The membrane is washed three times with TBS-T and then treated with HRP-conjugated secondary antibody (MBL, 458, 1:3,000) in TBS-T supplemented with 1% nonfat dry milk for 1 h at room temperature. The membrane is washed three times with TBS-T. Chemiluminescent signals generated with ECL Start or ECL Prime (GE Healthcare) were detected with a Fusion Solo S imaging system (Vilber Lourmat). The band intensity of GluA1 and GluA2 was normalized by the intensity of βIII-tubulin.

### Two-step labeling and PFQX1(AF488) staining in cultured neurons

To label endogenous AMPARs, 10 µM CAM2(TCO) in 200 µl of the culture medium was gently added to the hippocampal neurons cultured in 800 µl medium on a glass bottom dish to a final concentration of 2 µM CAM2(TCO). The cells were incubated for 2 h at 37 °C. In the second step, the culture medium was removed, and the cells were treated with 100 nM Tz(ST647) for 5 min in HEPES-based ACSF at room temperature. To quench excess Tz(ST647), 1 µM TCO-OH in HEPES-based ACSF was added. Then, the cells were treated with 100 nM PFQX1(AF488). Imaging was performed with confocal microscopy.

### Statistical analysis

All data are expressed as mean ± s.e.m. We accumulated the data for each condition from at least three independent experiments. We evaluated statistical significance with Student’s t-test or one-way ANOVA with Dunnet’s test. A value of *P* < 0.05 was considered significant.

## Data availability

Coordinates and structure factors have been deposited in the RCSB Protein Data Bank (PDB) with an accession code 9J8K for the S1S2J bound to PFQX1(AF488).

## Acknowledgements

We thank Eric Gouaux (Oregon Health and Science University) for the S1S2J construct. We also thank Mr. Yutaro Sugihara (Nagoya University) for the synthesis of Halo-tag probe, Mr. Jun Ikeshita (Nagoya University) for the construction of Halo-tag plasmid, Prof. Makoto Kinoshita (Nagoya University) for technical advice for the live imaging in neurons, and Dr. Bronwen Gardner from Edanz (https://jp.edanz.com/ac) for editing a draft of this manuscript. We thank the beamline scientists at SPring-8 BL45XU for supporting measurements. Data collections were carried out with the approval of the Japan Synchrotron Radiation Research Institute (Proposal number 2023B2746). This work was funded by Grants-in-Aid for Scientific Research (KAKENHI) (Grant Number 22KJ1605 to K.S., 24K02212 and 22K19364 to W.K., 23H05405 to I.H., 19H05781, 24H02262 and 24H00547 to E.N., 24H02265, 24H00492 and 24K21823 to S.K.), Novartis Foundation (to S.K.), the Takeda Science Foundation (to W.K and S.K.) and supported by JST CREST (JPMJCR1854) to M.Y. and JST ERATO (JPMJER1802) to I.H.

## Author contributions

S.K. initiated and designed the project. K.S. and M.N. performed the construction of AMPARs plasmids. K.S., M.N. and A.S. performed synthesis. H.A., T.F. and E.N. performed X-ray structural analysis. K.S. and S.K. performed confocal imaging in HEK293 cells and cultured neurons, and western blotting. W.K. and M.Y provided interpretation about results of synaptic plasticity. K.S., T.F., E.N. and S.K. wrote the manuscript. K.Y., W.K. and I.H. reviewed and edited the manuscript. All authors discussed and commented on the manuscript.

## Extended Data Figure

**Extended Data Fig. 1.**
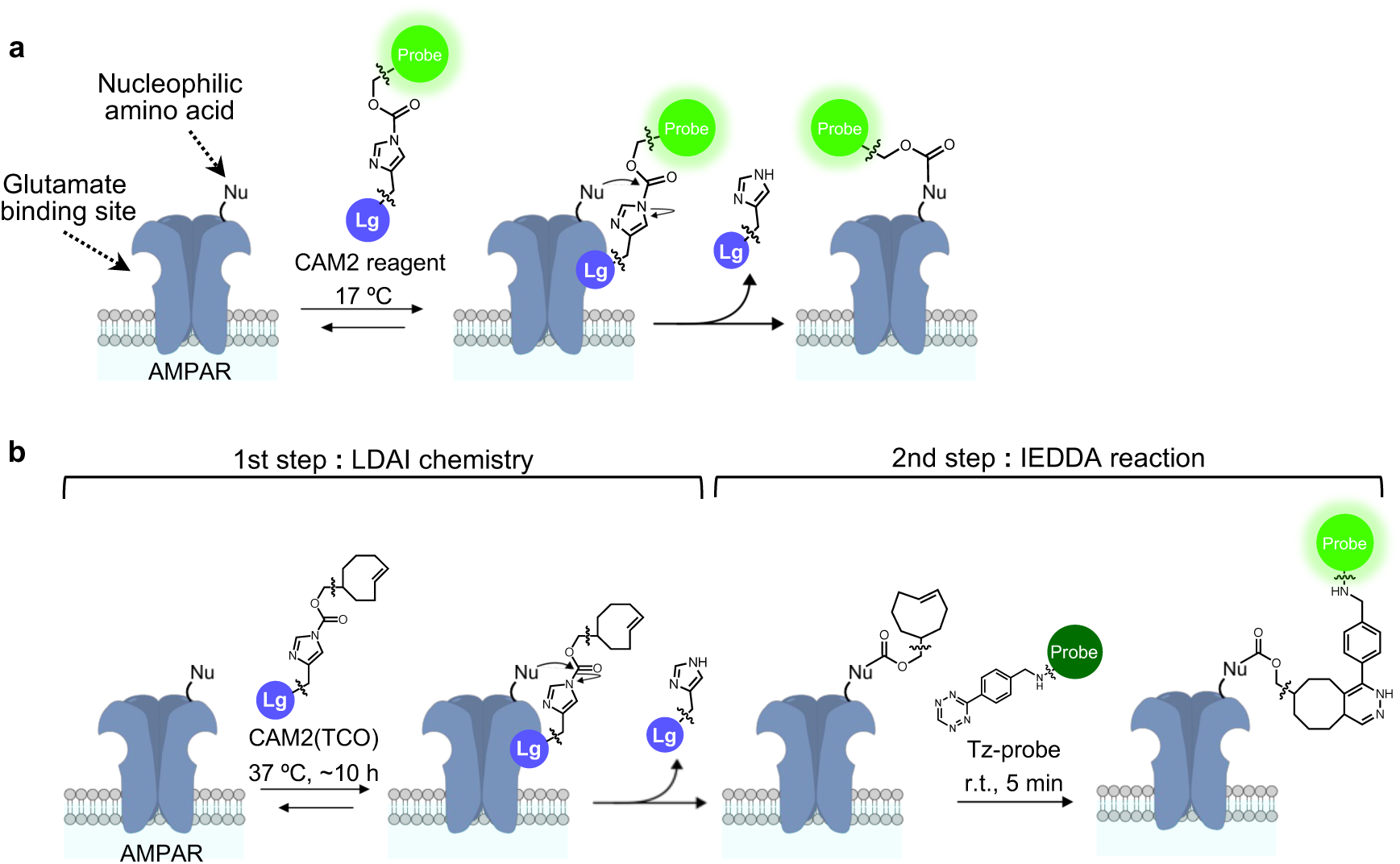
Ligand-directed chemical labeling of AMPARs. **(a)** A schematic illustration of ligand-directed acyl imidazole (LDAI) chemistry. CAM2 reagent is covalently attached to nucleophilic amino acid residues near the ligand binding site of AMPARs. To suppress the internalization of AMPARs, the cells should be incubated with CAM2 reagent at 17 °C. **(b)** A schematic illustration of ligand-directed two-step labeling. In the 1st step, TCO is covalently attached to AMPARs using CAM2(TCO). In the 2nd step, tetrazine conjugated fluorescent dye (Tz-probe) is selectively tethered to TCO via inverse electron demand Diels-Alder (IEDDA) reaction. Cell-surface AMPARs can be selectively labeled with the dye because of the extremely high reaction rate of IEDDA reaction.

**Extended Data Fig. 2.**
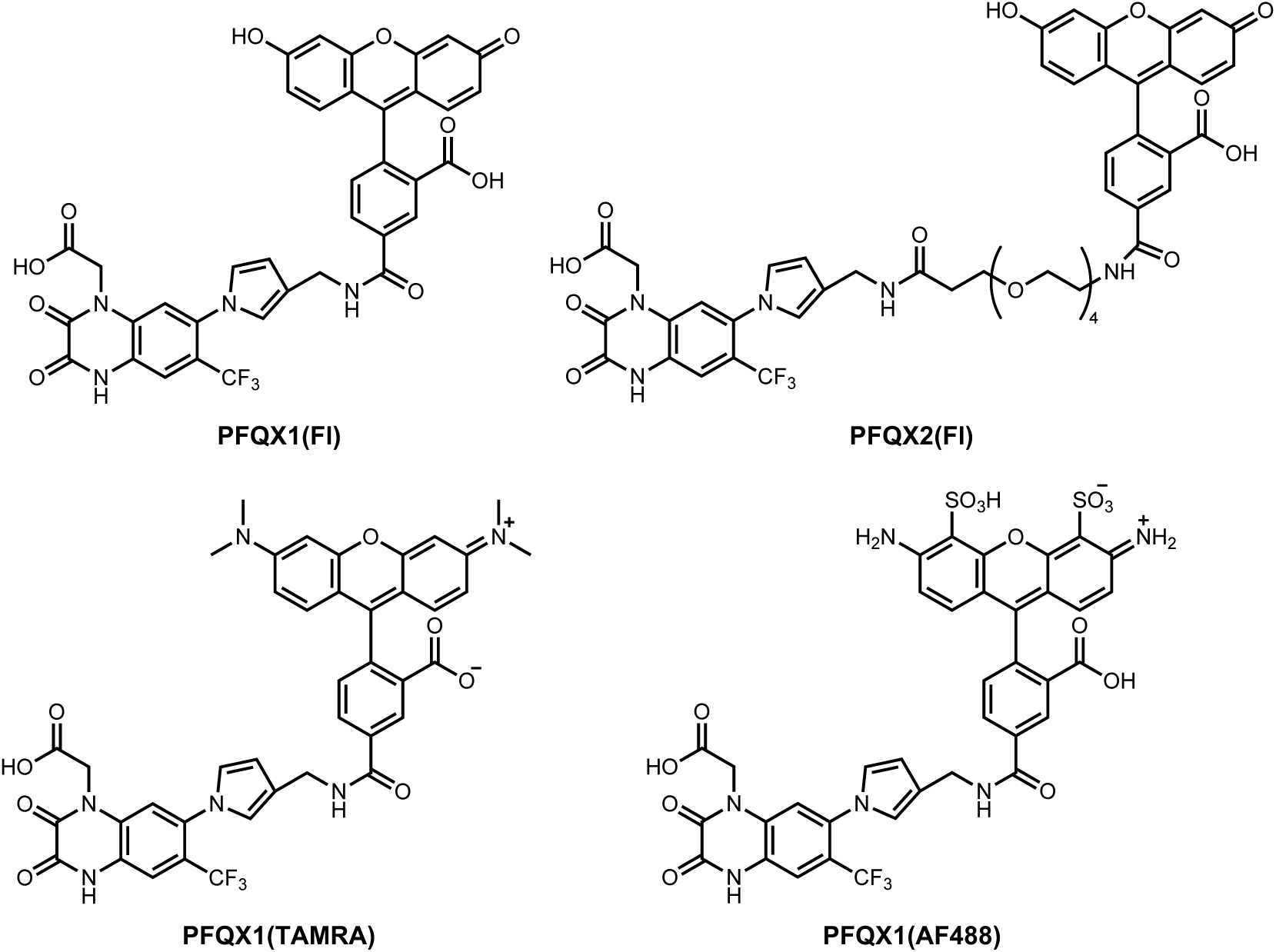
Detailed chemical structures of fluorophore–ligand conjugates. Chemical structures of PFQX1(Fl), PFQX2(Fl), PFQX1(TAMRA) and PFQX1(AF488) are shown.

**Extended Data Fig. 3.**
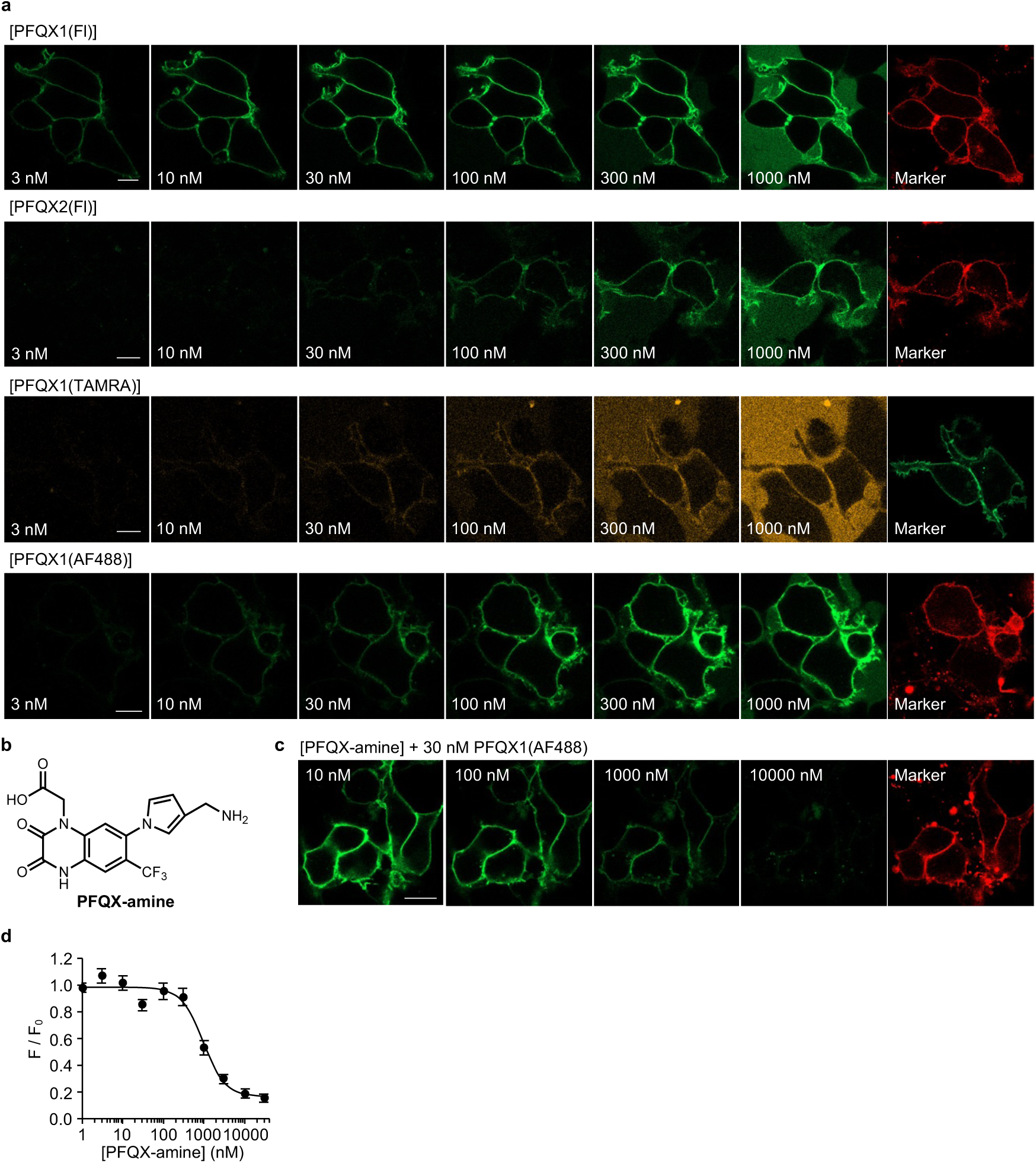
Concentration-dependency of fluorophore–ligand conjugates and the affinity of PFQX-amine to AMPARs. **(a)** Concentration-dependent binding of fluorophore–ligand conjugates to GluA2. Representative results of confocal live cell imaging of HEK293T cells transfected with GluA2 are shown. mCherry-F (for Fl and AF488) and EGFP-F (for TAMRA) were used as transfection markers. Scale bars, 10 µm. **(b–d)** Competitive inhibition of PFQX1(AF488) binding by PFQX-amine. In **b**, chemical structure of PFQX-amine, the synthetic intermediate for fluorophore–ligand conjugates is shown. In **c**, representative confocal live imaging of HEK293T cells transfected with GluA2 in the presence of each concentration of PFQX-amine are shown. In **d**, concentration-dependency of PFQX-amine is shown. The surface intensity of PFQX1(AF488) was quantified, which was normalized by that of 0 nM PFQX-amine treatment. [PFQX1(AF488)] = 30 nM (n = 3). Data are represented as mean ± s.e.m.

**Extended Data Fig. 4.**
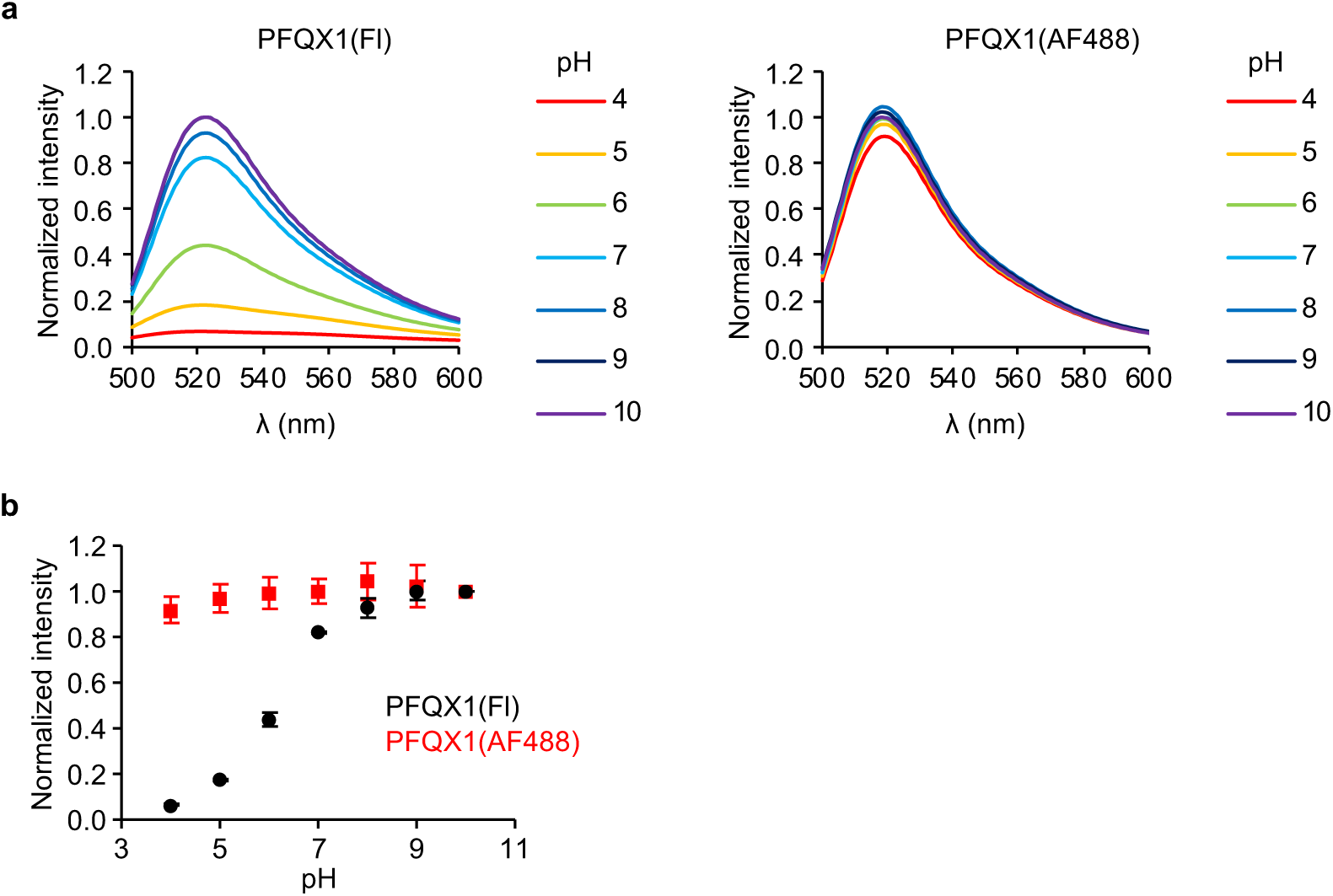
pH-dependency of PFQX1(Fl) and PFQX1(AF488). **(a)** Fluorescence spectra of PFQX1(Fl) (left) and PFQX1(AF488) (right) are shown. The fluorescence intensity was normalized by that at pH 10. Samples were prepared in GTA buffer (pH4–10). **(b)** pH titration curve of PFQX1(Fl) and PFQX1(AF488). [PFQX1(Fl)] = 2 µM, [PFQX1(AF488)] = 2 µM (n = 3). λ_ex_ = 480 nm. Data are represented as mean ± s.e.m.

**Extended Data Fig. 5.**
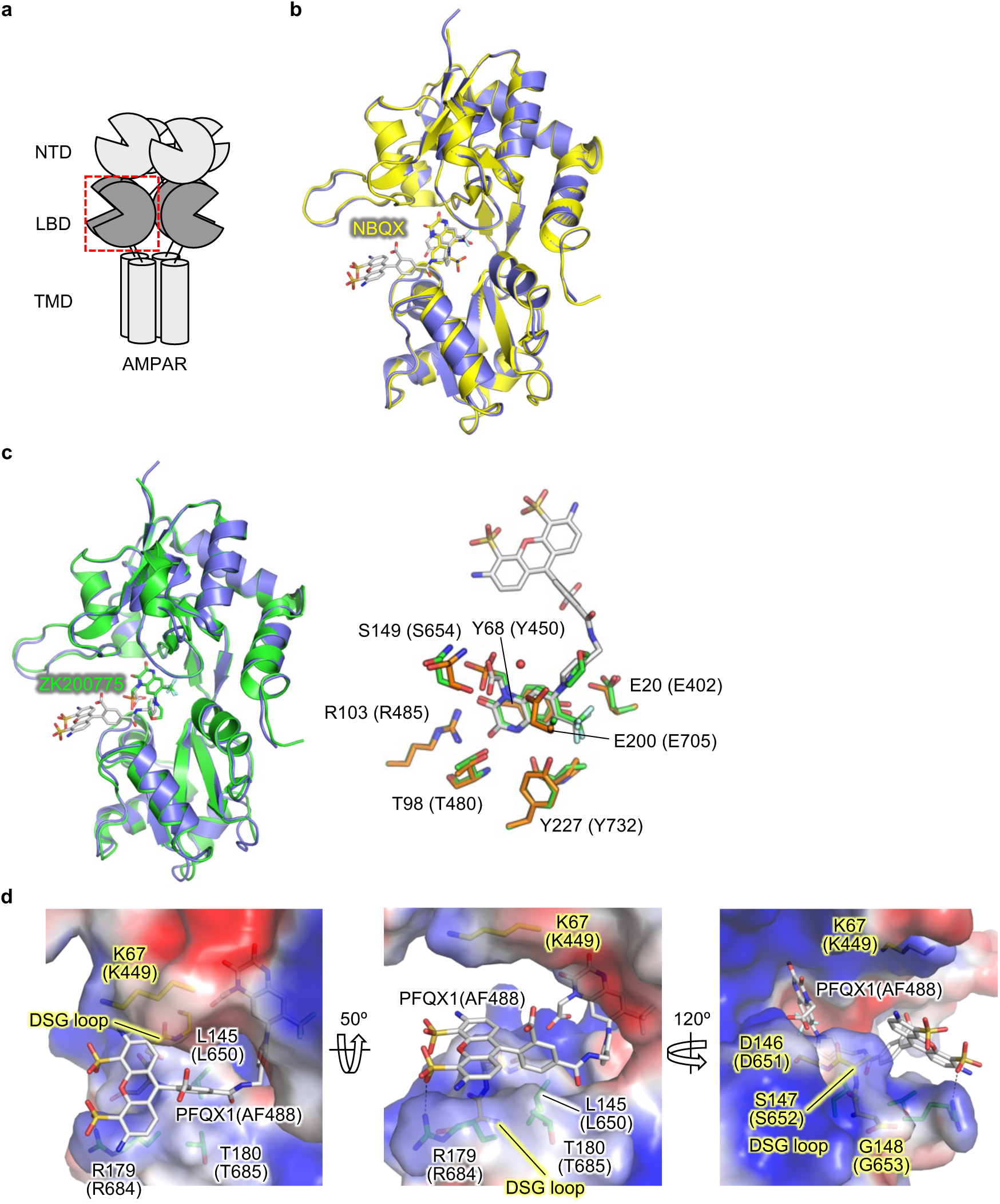
Comparison of the X-ray structure of PFQX1(AF488) bound S1S2J with other antagonist bound forms. **(a)** Schematic representation of AMPAR. Each subunit of AMPAR is composed of the amino-terminal domain (NTD), the ligand-binding domain (LBD), and the transmembrane domain (TMD). **(b)** Superposed structures of S1S2J bound to NBQX (PDB code: 6FQH) (yellow) and PFQX1(AF488) (purple). Bound NBQX and PFQX1(AF488) are shown as sticks. **(c)** Superposed structures of ligand binding domain bound to ZK200757 (PDB code: 5KBV) (green) and PFQX1(AF488) (purple). Bound ZK200757 and PFQX1(AF488) are shown as sticks. Right, close-up view of the ligand binding site. ZK200757 and the residues interacted with the ligand are shown as sticks (green). PFQX1(AF488) and the residues interacted with PFQX part of the ligand are also shown as sticks (orange and white, respectively). **(d)** The electrostatic potential around the AF488 binding region of the S1S2J. The residues involved in recognizing the AF488 part are shown as sticks (green), while Lys67 (Lys449) and DSG loop are also shown as sticks (yellow).

**Extended Data Fig. 6.**
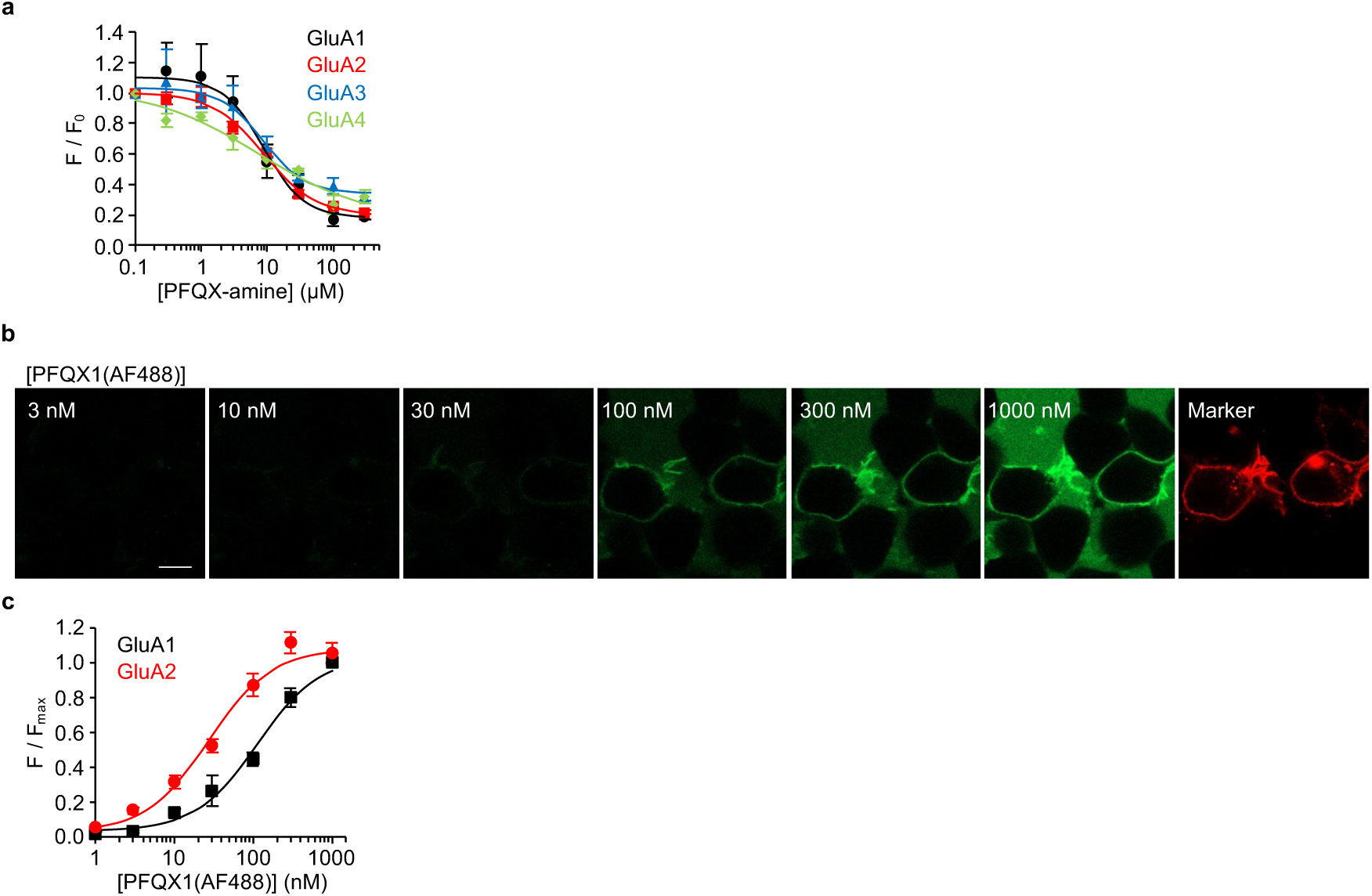
Non-specific action of PFQX derivatives to AMPAR subunits. **(a)** Fluorescence Ca^2+^ imaging assay to analyze inhibition of 30 µM glutamate-induced response by PFQX-amine. HEK293T cells were transfected with GluA1, GluA2, GluA3 (Y454A/R461G), or GluA4. Mutation was introduced to GluA3 to enhance surface expression of the homomer. **(b)** Representative results of confocal live cell imaging of HEK293T cells expressed with GluA1 are shown. The cells were treated with PFQX1(AF488) at each concentration. Scale bar, 10 µm. **(c)** Surface intensity of PFQX1(AF488) was quantified (n = 3). Data are represented as mean ± s.e.m.

**Extended Data Fig. 7.**
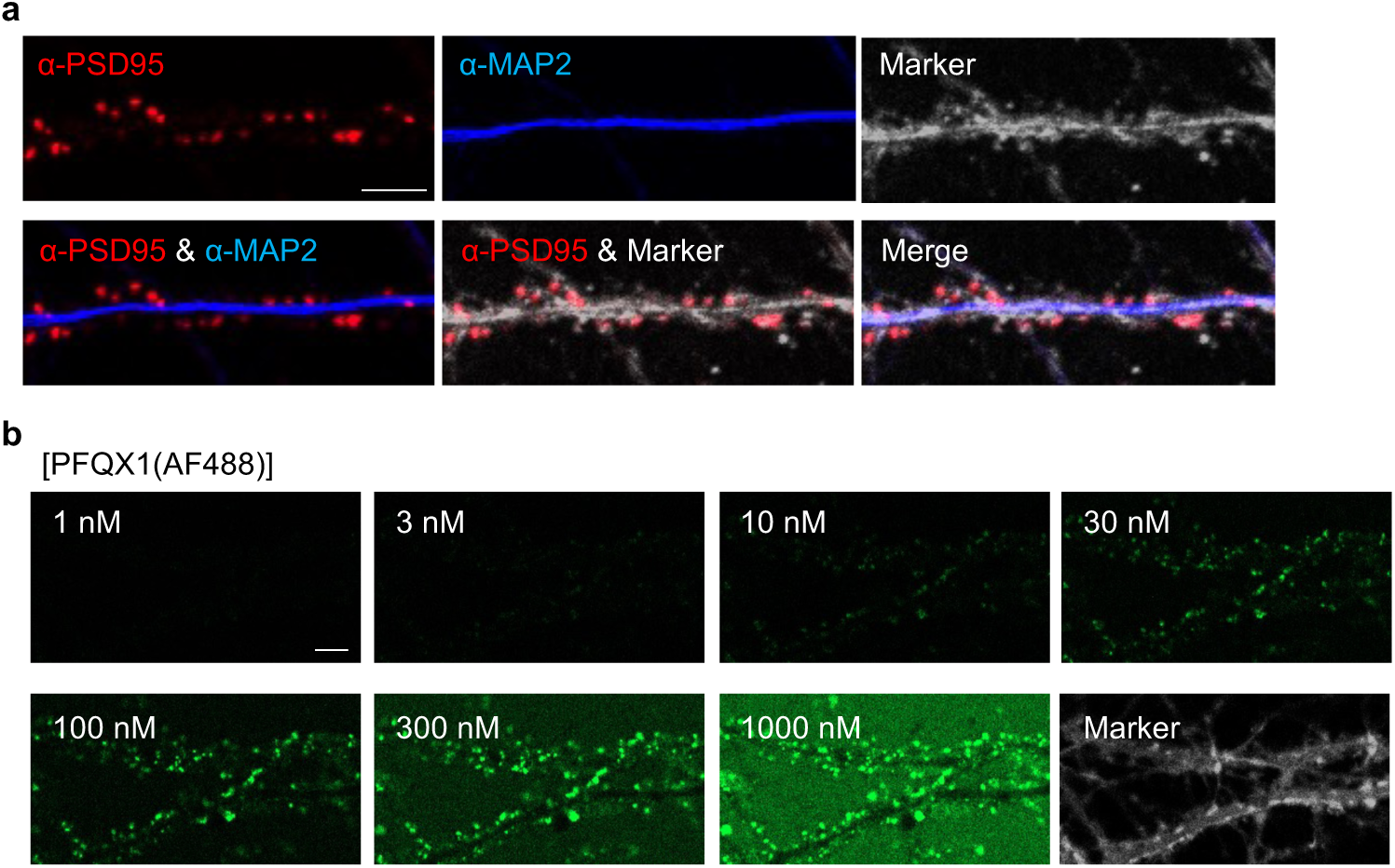
Immunostaining and concentration-dependency of PFQX-1(AF488) binding to AMPARs in cultured hippocampal neurons. **(a)** Immunostaining of cultured hippocampal neurons. The cells were immunostained with anti-PSD95 (red signal) and anti-MAP2 (blue signal) after staining with CellTracker Red CMPTX (gray signal). Scale bar, 5 µm. **(b)** Concentration-dependency of PFQX1(AF488) binding to AMPARs in cultured hippocampal neurons. Representative results of confocal live cell imaging are shown. Scale bar, 5 µm.

**Extended Data Fig. 8.**
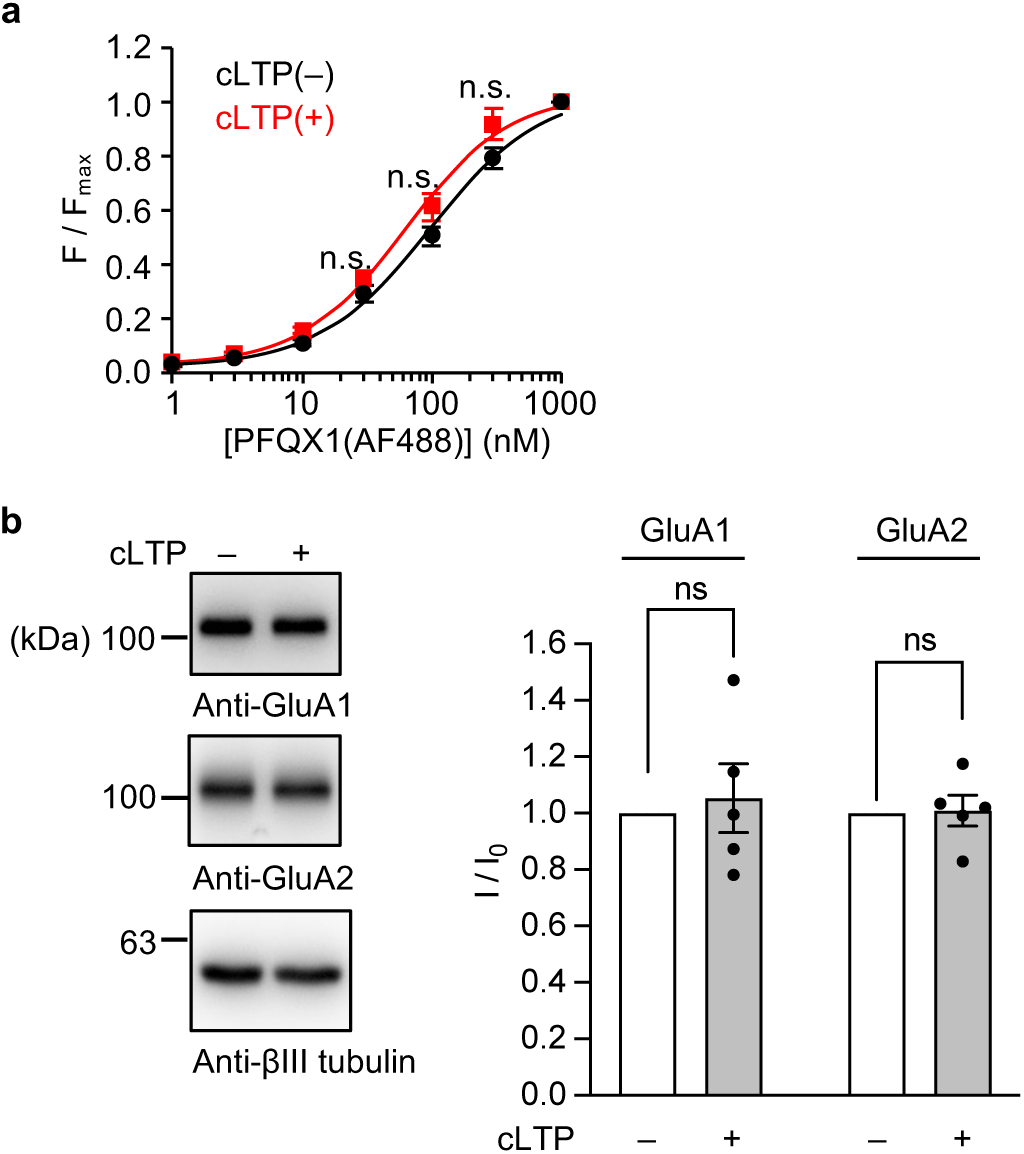
Binding affinity to PFQX1(AF488) and expression level of AMPARs were not affected by cLTP. **(a)** Concentration-dependency of PFQX1(AF488) binding to AMPARs in cultured hippocampal neurons was evaluated before and after cLTP (n = 3). **(b)** Western blotting analysis of cultured hippocampal neurons. The expression levels of GluA1, GluA2, and βIII tubulin as a loading control were analyzed before and after cLTP stimulation. Left, representative blots are shown. Right, quantification of band intensity. Band intensity was normalized by cLTP(–) (n = 5). ns, not significant (p > 0.05, Student’s t-test.). Data are represented as mean ± s.e.m.

**Extended Data Fig. 9.**
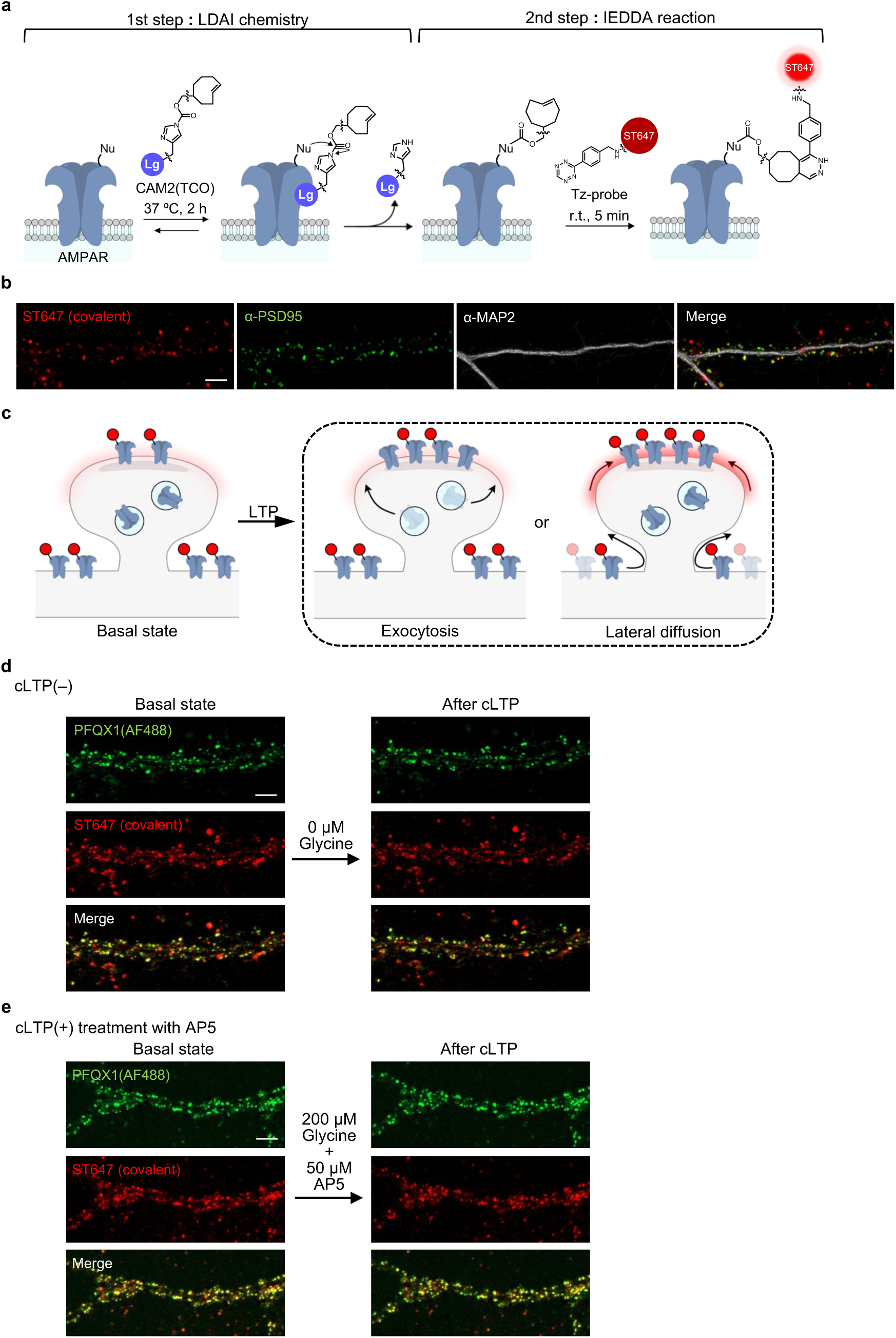
Covalent ST647 labeling by ligand-directed two-step labeling and evaluation of fluorescent changes after cLTP stimulation. **(a)** Schematic illustration of ligand-directed two-step labeling method. In the first step, a strained alkene (TCO) is covalently attached to AMPARs. In the second step, Tz(ST647) is selectively tethered to TCO via IEDDA reaction. **(b)** Immunostaining of cultured hippocampal neurons after labeling with ST647. The cells were immunostained with anti-PSD95 (green signal) and anti-MAP2 (gray signal). [CAM2(TCO)] = 2 µM, [Tz(ST647)] = 100 nM. Scale bar, 5 µm. **(c)** A schematic illustration of putative results of AMPAR trafficking after cLTP stimulation, where cell-surface AMPARs are covalently labeled with ST647 by two-step labeling. If exocytosis was the main mechanism of synaptic accumulation, non-labeled AMPARs would be exocytosed, resulting in no change in the intensity of ST647 in spines. If lateral diffusion was the main mechanism, labeled AMPARs would be accumulated, resulting in an enhanced intensity of ST647 in spines. **(d, e)** Representative live cell images of PFQX1(AF488) for cLTP stimulation under the control conditions. In **d**, glycine was not added to the cells. In **e**, AP5 was pre-treated before cLTP stimulation. The experimental procedure and analyzed data are shown in Fig. 6c and 6e, respectively. Scale bar, 5 µm.

**Extended Data Fig. 10.**
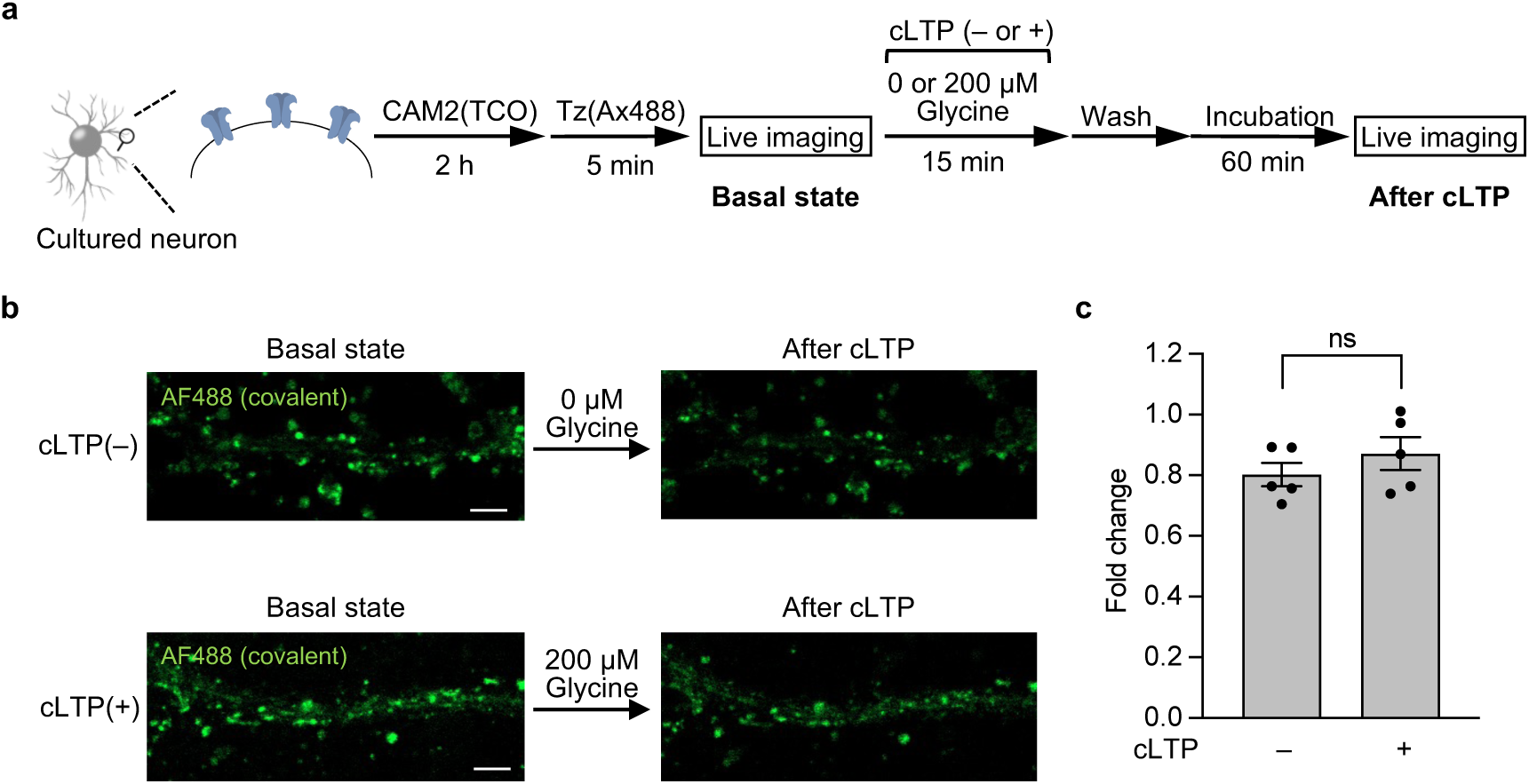
The fluorescent dye had no effect on the trafficking of AMPARs. **(a)** Schematic illustration of the experimental procedure. Cultured hippocampal neurons were labeled with 2 µM CAM2(TCO) for 2 h followed by Tz(Ax488) for 5 min. The cells were scanned before cLTP as “basal state” images. The cells were treated with 200 µM glycine for 15 min, and further incubated for 60 min. Then, the cells were scanned as “after cLTP” images. **(b)** Representative images of confocal live cell imaging covalently modified with Ax488 are shown. Upper, images of the control condition where glycine was not added to the cells are shown. Lower, images of the cells stimulated with glycine are shown. Scale bars, 5 µm. **(c)** The intensity of Ax488 in spines was analyzed, with the intensity of “after cLTP” images divided by that of “basal state” images (n = 5). ns, not significant (p > 0.05, Student’s t-test.). Data are represented as mean ± s.e.m.

## Supplementary Information

**Supplementary Table 1.** Data collection and refinement statistics.

|  | S1S2J-PFQX1(AF488) |
| --- | --- |
| **Data collection** |  |
| Wavelength | 1.000 |
| Resolution range (Å) | 47.33 – 1.82 (1.89 – 1.82) |
| Space group | *P*6<sub>1</sub>22 |
| Unit cell (*a*, *b*, *c*, $\alpha$, $\beta$, $\gamma$) (Å, °) | 69.9, 69.9, 227.5, 90, 90, 120 |
| Unique reflections | 30612 (2985) |
| Multiplicity | 71.2 (44.4) |
| Completeness (%) | 100.0 (100.0) |
| Mean *I*/σ*I* | 40.92 (5.05) |
| *R*<sub>merge</sub><sup>a</sup> | 0.181 (3.369) |
| *CC*<sub>1/2</sub> | 1.000 (0.983) |
| **Refinement** |  |
| Reflections for refinement | 30604 |
| *R*<sub>work</sub><sup>b</sup>/*R*<sub>free</sub><sup>c</sup> | 0.169 / 0.200 |
| RMSDs |  |
| lengths (Å) | 0.010 |
| angles (°) | 1.078 |
| Ramachandran (%) |  |
| favored | 99.22 |
| allowed | 0.78 |
| outliers | 0 |
| Rotamer outliers (%) | 0 |
| C–beta outliers (%) | 0 |
| Number of non-hydrogen atoms |  |
| S1S2J | 2040 |
| waters | 237 |
| PFQX1(AF488) | 62 |
| sulfates | 10 |
| polyethylene glycols | 14 |
| Mean B value (Å<sup>2</sup>) |  |
| overall | 33.92 |
| S1S2J | 32.61 |
| waters | 40.50 |
| PFQX1(AF488) | 39.79 |
| sulfates | 71.58 |
| polyethylene glycols | 60.24 |

<sup>a</sup>*R*<sub>merge</sub> = $\sum hkl \sum i |I_i(hkl) - \langle I_i(hkl) \rangle| / \sum hkl \sum i I_i(hkl)$ , where *i* is the number of observations of a given reflection and *I*(*hkl*) is the average intensity of the *i* observations. <sup>b</sup>*R*<sub>work</sub> = $\sum hkl ||F_{\text{obs}}(hkl)| - |F_{\text{calc}}(hkl)|| / \sum hkl |F_{\text{obs}}(hkl)|$ . <sup>c</sup>*R*<sub>free</sub> was calculated with a 5% fraction of randomly selected reflections evaluated from refinement. The highest resolution shell is shown in parentheses.

**Supplementary Table 2.** Interactions between PFQX1(AF488) and S1S2J.

| residues | atom type | ligand | atoms<sup>a</sup> | distance | interaction |
| --- | --- | --- | --- | --- | --- |
| E20 (E402) | Cγ | PFQX | F1 | 3.1 | multipolar F |
| Y68 (Y450) | phenyl | PFQX | quinoxalin | 3.6 | π-stacking |
| Y68 (Y450) | phenyl | PFQX | quinoxalin | 4.1 | π-stacking |
| P96 (P478) | O (main | PFQX | N4 | 2.8 | H-bond |
| T98 (T480) | N (main | PFQX | O2 | 2.8 | H-bond |
| R103 (R485) | Nη1 | PFQX | O2 | 2.7 | H-bond |
| R103 (R485) | Nη2 | PFQX | O3 | 2.9 | H-bond |
| S149 (S654) | N (main | PFQX | O5 | 2.9 | H-bond |
| S149 (S654) | Oγ | PFQX | O4 | 2.7 | H-bond |
| Y227 (Y732) | OH | PFQX | F2 | 3.1 | H-bond |
| water122 | O | PFQX | O5 | 3.1 | H-bond |
| water186 | O | PFQX | O4 | 2.6 | H-bond |
| water69 | O | PFQX | N2 | 3.2 | H-bond |
| T181 (T686) | N (main | AF48 | O1 | 2.8 | H-bond |
| T181 (T686) | Oγ | AF48 | O1 | 2.7 | H-bond |
| L145 (L650) | Cδ | AF48 | C8 | 3.9 | hydrophobic |
| R179 (R684) | Nη1 | AF48 | O8 | 2.7 | salt bridge |
| T180 (T685) | methyl | AF48 | phenyl | 4.2 | CH-π interaction |
| water113 | O | AF48 | O6 | 2.7 | H-bond |
| symmetry |  |  |  |  |  |
| K151 (K656) | Nζ | AF48 | O8 | 3.0 | salt bridge |
| K151 (K656) | Nζ | AF48 | O9 | 3.4 | salt bridge |
| K151 (K656) | Nζ | AF48 | O12 | 3.3 | salt bridge |
| K151 (K656) | Nζ | AF48 | O13 | 3.0 | salt bridge |
| R155 (R660) | Nη2 | AF48 | O12 | 3.2 | salt bridge |
| R155 (R660) | Nη2 | AF48 | O14 | 3.0 | salt bridge |
| R170 (R675) | Nη1 | AF48 | O9 | 3.2 | salt bridge |
| R170 (R675) | Nη1 | AF48 | O12 | 3.4 | salt bridge |
| R170 (R675) | O (main | AF48 | N6 | 3.0 | H-bond |
| R170 (R675) | guanidino | AF48 | xanthene | 3.6 | cation-π |

<sup>a</sup>Atomic number of PFQX1(AF488) is described in the below structure.

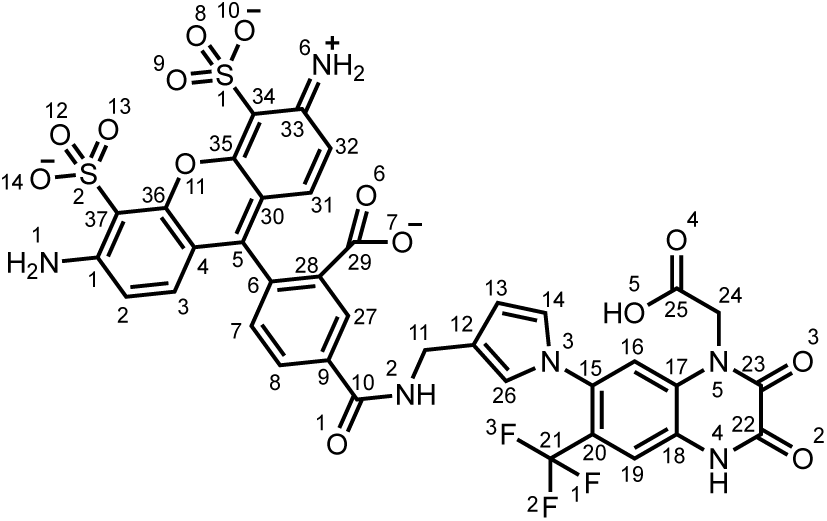

### Synthesis and Characterization

#### General materials and methods for organic synthesis

All chemical reagents and solvents were purchased from commercial sources (FUJIFILM Wako pure chemical, TCI chemical, Sigma-Aldrich, Click Chemistry Tools, BroadPharm) and were used without further purification. Thin-layer chromatography (TLC) was performed on silica gel 60 F254 precoated aluminum sheets (Merck). Chromatographic purification was performed by flash column chromatography on silica gel 60 N (neutral, 40–50 µm, Kanto Chemical). ^1^H-NMR spectra were recorded in deuterated solvents on an UltraShield 300 or 500 (Bruker). Chemical shifts were referenced to residual solvent peaks or tetramethyl silane (δ = 0 ppm). Multiplicities are abbreviated as follows: s = singlet, d = doublet, dd = double boublet, t = triplet, m = multiplet. High-resolution mass spectra were measured on a Compact (Bruker) equipped with electron spray ionization (ESI). Reversed-phase HPLC (RP-HPLC) was performed on a Hitachi Chromaster system equipped with a diode array and a YMC-Pack ODS-A column.

##### Synthesis of PFQX1(Fl)

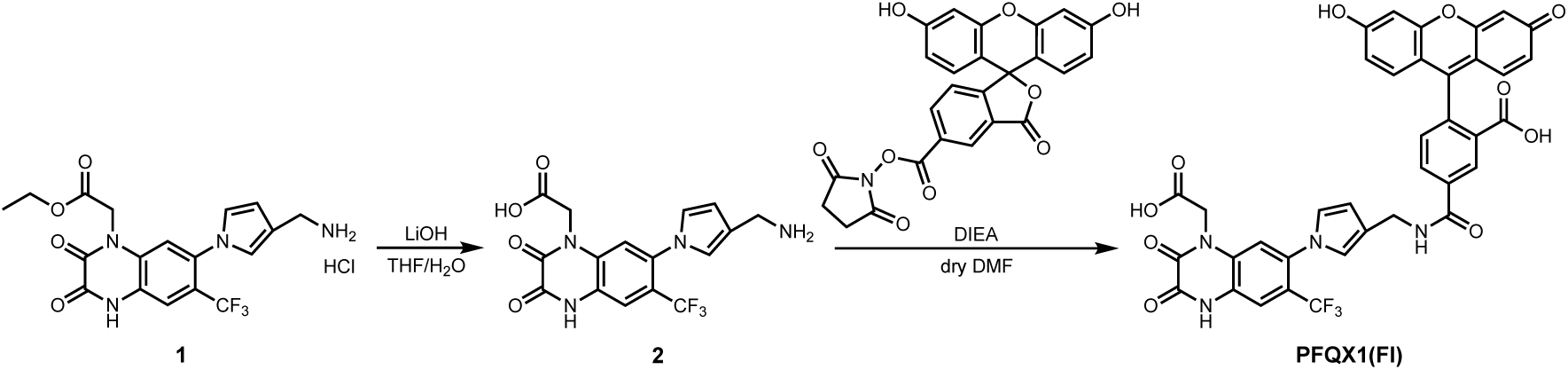

Synthesis of compound **2 (PFQX-amine)**

A solution of **1**^S1^ (64 mg, 0.14 mmol) and LiOH·H_2_O (12 mg, 0.28 mmol) in THF (0.5 mL) and H_2_O (0.5 mL) was stirred for 16 h at room temperature. After neutralization of the solvent with 1 M HCl, the crude was passed through a short pad of silica gel to give white crude **2** (76 mg) and used for the next step without further purification. A small portion of the crude was purified by RP-HPLC for the biological assay. The HPLC condition was as follows; (ODS-A, 250 × 20 mm, mobile phase; CH_3_CN (containing 0.1% TFA) : H_2_O (containing 0.1% TFA) = 10 : 90 → 30 : 70 (linear gradient over 20 min), flow rate; 10 mL/min, detection; UV (250 nm)). ^1^H-NMR (300 MHz, CD_3_OD) δ 7.52 (s, 1H), 7.13 (s, 1H), 6.94 (s, 1H), 6.82 (s, 1H), 6.30-6.28 (m, 1H), 4.87 (s, 2H), 3.94 (s, 2H). HR-ESI MS m/z calcd for [M+Na]^+^ 405.0781, found 405.0762.

##### Synthesis of **PFQX1(Fl)**

A solution of crude **2** (1.8 mg), DIEA (4.4 µL, 25 µmol), and 5-carboxyfluorescein NHS ester (1.0 mg, 2.1 µmol) in dry DMF (1 mL) was stirred for 15.5 h at room temperature under a nitrogen atmosphere. The crude mixture was purified by RP-HPLC (ODS-A, 250 × 10 mm, mobile phase; CH_3_CN (containing 0.1% TFA) : H_2_O (containing 0.1% TFA) = 10 : 90 → 50 : 50 (linear gradient over 40 min), flow rate; 3 mL/min, detection; UV (250 nm)) to give **PFQX1(Fl)** (0.97 mg, 1.3 µmol, 62% yield) as an orange solid. ^1^H-NMR (500 MHz, CD_3_OD) δ 8.54 (s, 1H), 8.25 (dd, *J* = 1.5, 8.5 Hz, 1H), 7.61 (s, 1H), 7.38 (d, *J* = 8.0 Hz, 1H), 7.29 (s, 1H), 6.92 (s, 1H), 6.86-6.83 (m, 5H), 6.71 (d, *J* = 8.0 Hz, 2H), 6.35 (s, 1H), 5.02 (s, 2H), 4.54 (s, 2H). HR-ESI MS m/z calcd for [M-H]^-^ 739.1294, found 739.1305.

##### Synthesis of PFQX2(Fl)

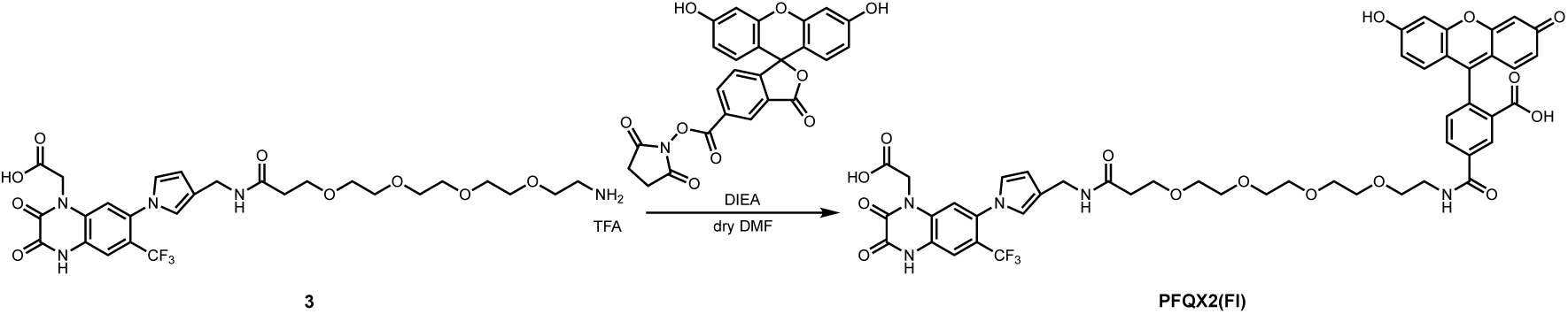

A solution of **3**^S1^ (4.0 mg), DIEA (10 µL, 57 µmol), and 5-carboxyfluorescein NHS ester (1.0 mg, 2.1 µmol) in dry DMF (1 mL) was stirred for 15.5 h at room temperature under a nitrogen atmosphere. The crude mixture was purified by RP-HPLC (ODS-A, 250 × 10 mm, mobile phase; CH_3_CN (containing 0.1% TFA) : H_2_O (containing 0.1% TFA) = 10 : 90 → 60 : 40 (linear gradient over 50 min), flow rate; 3 mL/min, detection; UV (250 nm)) to give **PFQX-L-Fl** (1.3 mg, 1.3 µmol, 62% yield) as an orange solid. δ 8.54 (s, 1H), 8.24 (dd, *J* = 1.5, 8.0 Hz, 1H), 7.59 (s, 1H), 7.37 (d, *J* = 8.0 Hz, 1H), 7.24 (s, 1H), 6.87-6.71 (m, 8H), 6.21 (s, 1H), 4.98 (s, 2H), 4.26 (s, 2H), 3.72-3.54 (m, 18 H), 2.45 (t, *J* = 6.0 Hz, 2H). HR-ESI MS m/z calcd for [M-H]^-^ 986.2713, found 986.2690.

##### Synthesis of PFQX1(TAMRA)

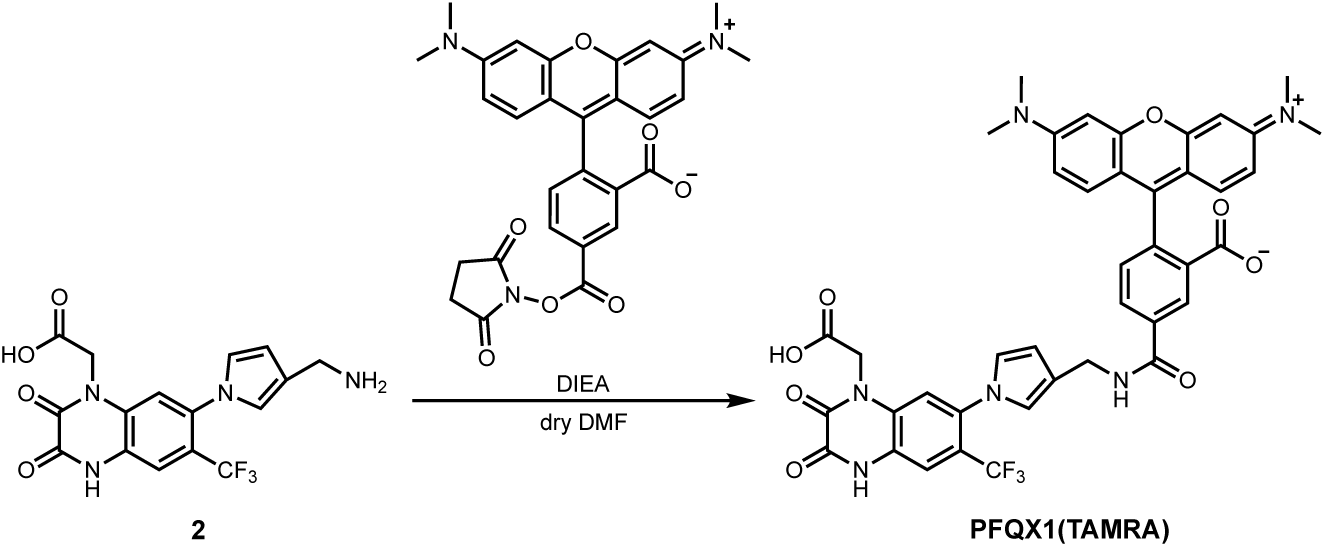

A solution of crude **2** (1.5 mg), DIEA (1.6 µL, 10 µmol), and 5-TAMRA NHS ester (1.0 mg, 1.9 µmol) in dry DMF (1 mL) was stirred for 16 h at room temperature under a nitrogen atmosphere. The crude mixture was purified by RP-HPLC (ODS-A, 250 × 10 mm, mobile phase; CH_3_CN : 10 mM AcONH_4_ aq. = 10 : 90 → 70 : 30 (linear gradient over 60 min), flow rate; 3 mL/min, detection; UV (250 nm)) to give **PFQX1(TAMRA)** (0.96 mg, 1.2 µmol, 63% yield) as a pink solid. ^1^H-NMR (300 MHz, CD_3_OD) δ 8.65 (s, 1H), 8.16 (dd, *J* = 7.8 Hz, 1H), 7.58 (s, 1H), 7.43 (d, *J* = 7.8 Hz, 1H), 7.22 (m, 3H), 7.05 (dd, *J* = 9.6, 2.4 Hz, 2H), 6.93 (m, 3H), 6.82 (s, 1H), 6.34 (t, *J* = 1.2 Hz, 1H), 4.86 (s, 2H), 4.54 (s, 2H), 3.29 (s, 12 H). HR-ESI MS m/z calcd for [M+H]^+^ 795.2385, found 795.2386.

##### Synthesis of PFQX1(AF488)

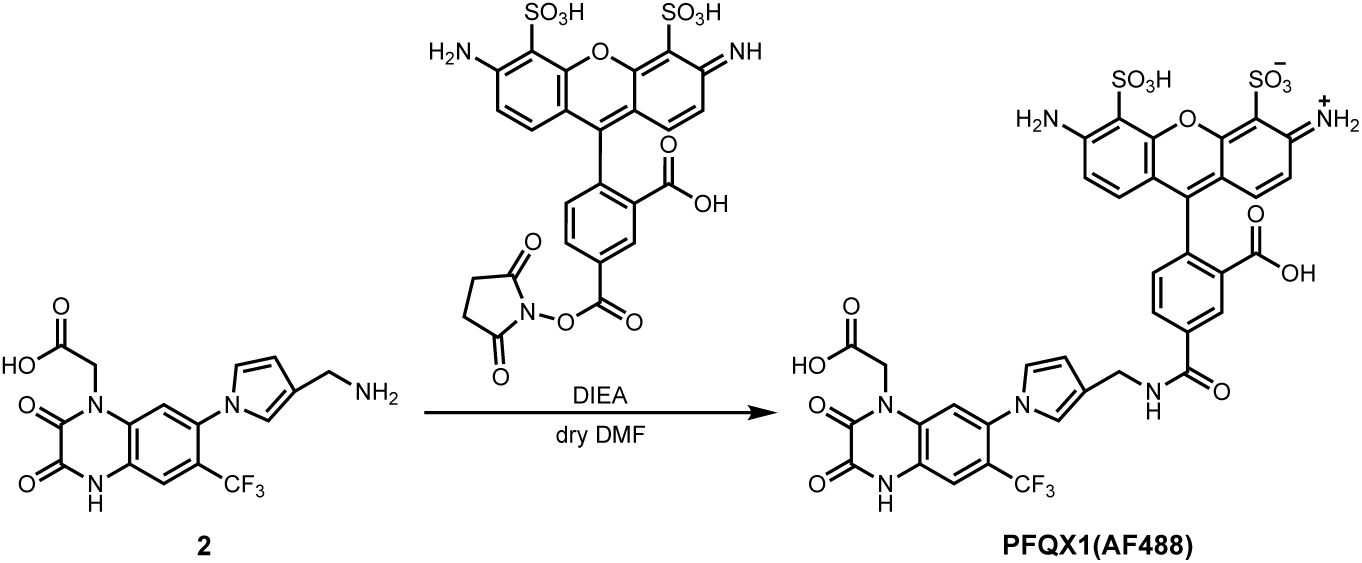

A solution of crude **2** (1.3 mg), DIEA (1.6 µL, 10 µmol), and Az488 NHS ester (1.0 mg, 1.6 µmol) in dry DMF (1 mL) was stirred for 16 h at room temperature under a nitrogen atmosphere. The crude mixture was purified by RP-HPLC (ODS-A, 250 × 10 mm, mobile phase; CH_3_CN : 10 mM AcONH_4_ aq. = 0 : 100 → 40 : 60 (linear gradient over 40 min), flow rate; 3 mL/min, detection; UV (250 nm)) to give **PFQX1(AF488)** (1.0 mg, 1.1 µmol, 69% yield) as an orange solid. ^1^H-NMR (300 MHz, D_2_O) δ 8.29 (s, 1H), 7.97 (dd, *J* = 1.4, 7.9 Hz, 1H), 7.64 (s, 1H), 7.34 (d, *J* = 8.0 Hz, 1H), 7.27 (s, 1H), 7.17 (d, *J* = 9.3 Hz, 2H), 7.01 (s, 1H), 6.94 (s, 1H), 6.90 (d, *J* = 9.3 Hz, 2H), 6.38 (t, *J* = 2.2 Hz, 1H), 4.73 (s, 2H), 4.52 (s, 2H). HR-ESI MS m/z calcd for [M-2H+Na]^-^ 919.0569, found 919.0588.

##### Synthesis of HTL(AF647)

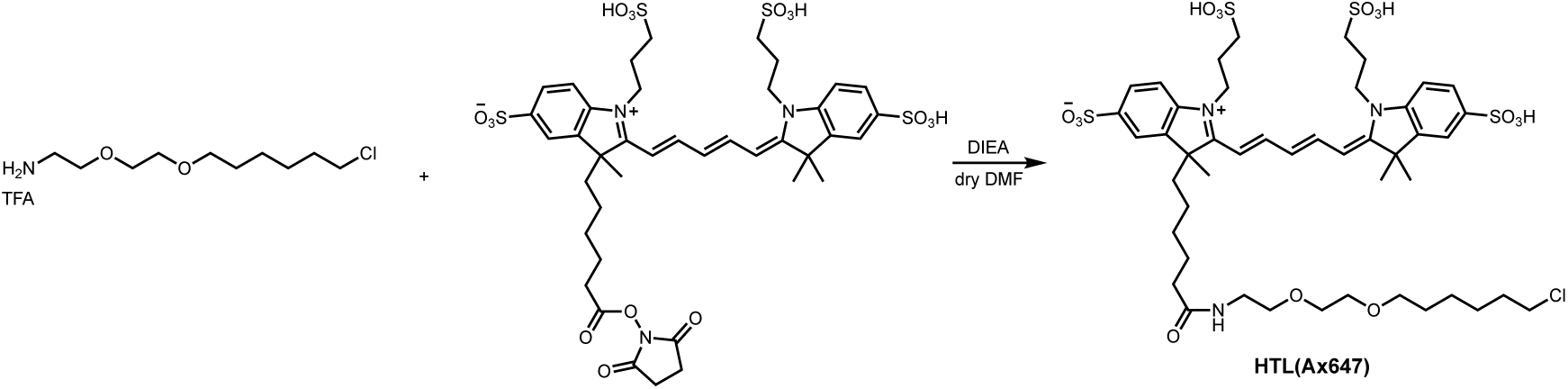

2-(2-((6-chlorohexyl)oxy)ethoxy) ethanamine TFA salt was synthesized as described in ref. S2. A solution of 2-(2-((6-chlorohexyl)oxy)ethoxy) ethanamine TFA salt (0.70 mg, 2.1 µmol), DIEA (1.0 µL, 6.0 µmol), and Alexa Fluor 647 NHS ester (1.0 mg, 1.0 µmol) in dry DMF (1 mL) was stirred for 19 h at room temperature under a nitrogen atmosphere. The crude mixture was purified by RP-HPLC (ODS-A, 250 × 10 mm, mobile phase; CH_3_CN : 10 mM AcONH_4_ aq. = 5 : 95 → 55 : 45 (linear gradient over 50 min), flow rate; 3 mL/min, detection; UV (250 nm)) to give **HTL(AF647)** (0.83 mg, 0.77 µmol, 74% yield) as a blue solid. HR-ESI MS m/z calcd for [M]•^+^ 1063.3060, found 1063.3067.

S1. Wakayama, S. *et al.* Chemical labelling for visualizing native AMPA receptors in live neurons. *Nat Commun* **8**, 14850 (2017).

S2. Los, G. V. *et al.* HaloTag: A Novel Protein Labeling Technology for Cell Imaging and Protein Analysis. *ACS Chem. Biol.* **3**, 373–382 (2008).

## References

1. Bliss, T. V. P. & Collingridge, G. L. A synaptic model of memory: long-term potentiation in the hippocampus. Nature 361, 31–39 (1993).

2. Martin, S. J., Grimwood, P. D. & Morris, R. G. M. Synaptic Plasticity and Memory: An Evaluation of the Hypothesis. Annu. Rev. Neurosci. 23, 649–711 (2000).

3. Citri, A. & Malenka, R. C. Synaptic Plasticity: Multiple Forms, Functions, and Mechanisms. Neuropsychopharmacology 33, 18–41 (2008).

4. Bredt, D. S. & Nicoll, R. A. AMPA Receptor Trafficking at Excitatory Synapses. Neuron 40, 361–379 (2003).

5. Huganir, R. L. & Nicoll, R. A. AMPARs and Synaptic Plasticity: The Last 25 Years. Neuron 80, 704–717 (2013).

6. Diering, G. H. & Huganir, R. L. The AMPA Receptor Code of Synaptic Plasticity. Neuron 100, 314–329 (2018).

7. Kopec, C. D., Li, B., Wei, W., Boehm, J. & Malinow, R. Glutamate Receptor Exocytosis and Spine Enlargement during Chemically Induced Long-Term Potentiation. J. Neurosci. 26, 2000–2009 (2006).

8. Tanaka, H. & Hirano, T. Visualization of Subunit-Specific Delivery of Glutamate Receptors to Postsynaptic Membrane during Hippocampal Long-Term Potentiation. Cell Rep. 1, 291–298 (2012).

9. Shepherd, J. D. & Huganir, R. L. The Cell Biology of Synaptic Plasticity: AMPA Receptor Trafficking. Annu. Rev. Cell Dev. Biol. 23, 613–643 (2007).

10. Ashby, M. C. et al. Removal of AMPA Receptors (AMPARs) from Synapses Is Preceded by Transient Endocytosis of Extrasynaptic AMPARs. J. Neurosci. 24, 5172–5176 (2004).

11. Makino, H. & Malinow, R. AMPA Receptor Incorporation into Synapses during LTP: The Role of Lateral Movement and Exocytosis. Neuron 64, 381–390 (2009).

12. Keppler, A. et al. A general method for the covalent labeling of fusion proteins with small molecules in vivo. Nat. Biotechnol. 21, 86–89 (2003).

13. Los, G. V. et al. HaloTag: A Novel Protein Labeling Technology for Cell Imaging and Protein Analysis. ACS Chem. Biol. 3, 373–382 (2008).

14. Harb, A. et al. Auxiliary Subunits Regulate the Dendritic Turnover of AMPA Receptors in Mouse Hippocampal Neurons. Front. Mol. Neurosci. 14, 728498 (2021).

15. Collingridge, G. L., Peineau, S., Howland, J. G. & Wang, Y. T. Long-term depression in the CNS. Nat. Rev. Neurosci. 11, 459–473 (2010).

16. Graves, A. R. et al. Visualizing synaptic plasticity in vivo by large-scale imaging of endogenous AMPA receptors. Elife 10, e66809 (2021).

17. Willems, J., et al. ORANGE: A CRISPR/Cas9-based genome editing toolbox for epitope tagging of endogenous proteins in neurons. PLoS Biol. 18, e3000665 (2020).

18. Fujishima, S.-H., Yasui, R., Miki, T., Ojida, A. & Hamachi, I. Ligand-Directed Acyl Imidazole Chemistry for Labeling of Membrane-Bound Proteins on Live Cells. J. Am. Chem. Soc. 134, 3961–3964 (2012).

19. Wakayama, S. et al. Chemical labelling for visualizing native AMPA receptors in live neurons. Nat. Commun. 8, 14850 (2017).

20. Nonaka, H. et al. Bioorthogonal chemical labeling of endogenous neurotransmitter receptors in living mouse brains. Proc. Natl. Acad. Sci. U. S. A. 121, e2313887121 (2024).

21. Ojima, K. et al. Ligand-directed two-step labeling to quantify neuronal glutamate receptor trafficking. Nat. Commun. 12, 831 (2021).

22. Lubisch, W., Behl, B. & Hofmann, H. P. Pyrrolylquinoxalinediones: A new class of AMPA receptor antagonists. Bioorg. Med. Chem. Lett. 6, 2887–2892 (1996).

23. Sobolevsky, A. I., Rosconi, M. P. & Gouaux, E. X-ray structure, symmetry and mechanism of an AMPA-subtype glutamate receptor. Nature 462, 745–756 (2009).

24. Traynelis, S. F. et al. Glutamate receptor ion channels: structure, regulation, and function. Pharmacol. Rev. 62, 405–496 (2010).

25. Schwenk, J. et al. Regional Diversity and Developmental Dynamics of the AMPA-Receptor Proteome in the Mammalian Brain. Neuron 84, 41–54 (2014).

26. Sinning, A. & Hübner, C. A. Minireview: pH and synaptic transmission. FEBS Lett. 587, 1923–1928 (2013).

27. Chen, G. Q., Sun, Y., Jin, R. & Gouaux, E. Probing the ligand binding domain of the GluR2 receptor by proteolysis and deletion mutagenesis defines domain boundaries and yields a crystallizable construct. Protein Sci. 7, 2623–2630 (1998).

28. Armstrong, N. & Gouaux, E. Mechanisms for activation and antagonism of an AMPA-sensitive glutamate receptor: crystal structures of the GluR2 ligand binding core. Neuron 28, 165–181 (2000).

29. Dürr, K. L. et al. Structure and dynamics of AMPA receptor GluA2 in resting, pre-open, and desensitized states. Cell 158, 778–792 (2014).

30. Coombs, I. D. et al. Homomeric GluA2(R) AMPA receptors can conduct when desensitized. Nat. Commun. 10, 4312 (2019).

31. Twomey, E. C., Yelshanskaya, M. V., Grassucci, R. A., Frank, J. & Sobolevsky, A. I. Elucidation of AMPA receptor-stargazin complexes by cryo-electron microscopy. Science 353, 83–86 (2016).

32. Lüscher, C. & Malenka, R. C. NMDA receptor-dependent long-term potentiation and long-term depression (LTP/LTD). Cold Spring Harb. Perspect. Biol. 4, a005710–a005710 (2012).

33. Bashir, Z. I., Alford, S., Davies, S. N., Randall, A. D. & Collingridge, G. L. Long-term potentiation of NMDA receptor-mediated synaptic transmission in the hippocampus. Nature 349, 156–158 (1991).

34. Lu, W.-Y. et al. Activation of Synaptic NMDA Receptors Induces Membrane Insertion of New AMPA Receptors and LTP in Cultured Hippocampal Neurons. Neuron 29, 243–254 (2001).

35. Zhang, X.-Y. et al. Glycine induces bidirectional modifications in N-methyl-D-aspartate receptor-mediated synaptic responses in hippocampal CA1 neurons. J. Biol. Chem. 289, 31200–31211 (2014).

36. Saneyoshi, T. et al. Reciprocal activation within a kinase-effector complex underlying persistence of structural LTP. Neuron 102, 1199–1210.e6 (2019).

37. Lisman, J., Yasuda, R. & Raghavachari, S. Mechanisms of CaMKII action in long-term potentiation. Nat. Rev. Neurosci. 13, 169–182 (2012).

38. Tan, H. L., Chiu, S.-L., Zhu, Q. & Huganir, R. L. GRIP1 regulates synaptic plasticity and learning and memory. Proc. Natl. Acad. Sci. U. S. A. 117, 25085–25091 (2020).

39. Kennedy, M. J., Davison, I. G., Robinson, C. G. & Ehlers, M. D. Syntaxin-4 Defines a Domain for Activity-Dependent Exocytosis in Dendritic Spines. Cell 141, 524– 535 (2010).

40. Esteves da Silva, M., et al. Positioning of AMPA Receptor-Containing Endosomes Regulates Synapse Architecture. Cell Rep. 13, 933–943 (2015).

41. Penn, A. C. et al. Hippocampal LTP and contextual learning require surface diffusion of AMPA receptors. Nature 549, 384–388 (2017).

42. Ashby, M. C., Maier, S. R., Nishimune, A. & Henley, J. M. Lateral diffusion drives constitutive exchange of AMPA receptors at dendritic spines and is regulated by spine morphology. J. Neurosci. 26, 7046–7055 (2006).

43. Rathje, M. et al. AMPA receptor pHluorin-GluA2 reports NMDA receptor-induced intracellular acidification in hippocampal neurons. Proc. Natl. Acad. Sci. U. S. A. 110, 14426–14431 (2013).

44. Studier, F. W., Rosenberg, A. H., Dunn, J. J. & Dubendorff, J. W. [6] Use of T7 RNA polymerase to direct expression of cloned genes. Methods in Enzymology 185, 60–89 (Academic Press, 1990).

45. Hirata, K. et al. ZOO: an automatic data-collection system for high-throughput structure analysis in protein microcrystallography. Acta Crystallogr. D Struct. Biol. 75, 138–150 (2019).

46. Liebschner, D. et al. Macromolecular structure determination using X-rays, neutrons and electrons: recent developments in Phenix. Acta Crystallogr. D Struct. Biol. 75, 861–877 (2019).

47. Emsley, P., Lohkamp, B., Scott, W. G. & Cowtan, K. Features and development of Coot. Acta Crystallogr. D Biol. Crystallogr. 66, 486–501 (2010).

48. Liebschner, D. et al. Polder maps: improving OMIT maps by excluding bulk solvent. Acta Crystallogr. D Struct. Biol. 73, 148–157 (2017).

49. Chen, V. B. et al. MolProbity: all-atom structure validation for macromolecular crystallography. Acta Crystallogr. D Biol. Crystallogr. 66, 12–21 (2010).

50. Kaech, S. & Banker, G. Culturing hippocampal neurons. Nat. Protoc. 1, 2406–2415 (2006).

